# Centrosome age breaks spindle size symmetry even in “symmetrically” dividing cells

**DOI:** 10.1101/2023.09.15.557935

**Authors:** Alexandre Thomas, Patrick Meraldi

## Abstract

Centrosomes are the main microtubule organizing center in animal cells. Due to the semi-conservative nature of centrosome duplication, the two centrosomes differ in age. In asymmetric stem cell divisions, centrosome age can induce an asymmetry in half-spindle lengths. However, whether centrosome age affects the symmetry of the two half-spindles in tissue culture cells thought to divide symmetrically, is unknown. Here, we show that in human epithelial and fibroblastic cell lines centrosome age imposes a subtle spindle asymmetry that leads to asymmetric cell daughter sizes. At the mechanistic level, we show that this asymmetry depends on the preferential accumulation on old centrosomes of the microtubule nucleation-organizing proteins pericentrin, γ-tubulin, Cdk5Rap2, and TPX2, under the control of a cenexin-bound pool of the mitotic kinase Plk1. Moreover, we find that old centrosomes have a higher microtubule nucleation capacity. We therefore postulate that centrosome age breaks spindle size symmetry via microtubule nucleation even in cells thought to divide symmetrically.

## INTRODUCTION

During mitosis, cells assemble a bipolar spindle to ensure faithful chromosome segregation between the two daughter cells (Prosser and Pelletier, 2017). This assembly relies on the dynamicity of spindle microtubules and the cooperative action of microtubule-associated proteins (MAPs) (Petry, 2016). In most animal cells, microtubules are first nucleated from the centrosome, the main microtubule organizing center (Sanchez and Feldman, 2017; Meraldi, 2016). Microtubules subsequently also emerge from chromosomes and kinetochores, the microtubule binding sites on chromosomes, and by branching off existing microtubule via the augmin complex (Heald et al., 1996; Gruss et al., 2002; Goshima et al., 2008; Uehara et al., 2009; Petry et al., 2013; David et al., 2019; Wu et al., 2023). Although centrosomes are dispensable for spindle assembly in various cell types (Heald et al., 1996; Basto et al., 2006; Bobinnec et al., 1998; Khodjakov et al., 2000), and absent in plants, planarians and in oocytes (Yi and Goshima, 2018; Mogessie et al., 2018; Azimzadeh et al., 2012), they regulate mitotic spindle assembly, and are crucial for spindle orientation and faithful chromosome segregation (Khodjakov and Rieder., 2001, (Khodjakov and Rieder, 2001; Buffin et al., 2007; Basto et al., 2006; Hayward et al., 2014; Sir et al., 2013; Dudka et al., 2019).

Centrosomes are composed of two orthogonally oriented centrioles surrounded by a pericentriolar material (PCM) required for microtubule nucleation (Conduit et al., 2015; Vasquez-Limeta and Loncarek, 2021). They duplicate once per cell cycle in a semi-conservative manner, as each existing centriole serves a seed for a new daughter centriole resulting in centrosomes of different ages (Nigg and Stearns, 2011; Nigg and Holland, 2018; Fırat-Karalar and Stearns, 2014). The duplicated old centrosome contains the oldest centriole called grandmother centriole and its daughter centriole, while the young centrosome contains a mother centriole and its daughter centriole (Sullenberger et al., 2020) The grandmother centriole is longer than the mother centriole and possesses distal (DA) and sub-distal (SDA) appendages, while mother centrioles only possess distal appendages. As cells enter mitosis, both centrosomes mature, massively expanding their PCM, which drives microtubule nucleation (Conduit et al., 2015; Palazzo et al., 2000). PCM expansion requires the activity of the mitotic kinase Plk1, which phosphorylates the pericentriolar proteins pericentrin and Cdk5Rap2/Cep215 (Lee and Rhee, 2011; Conduit et al., 2014; Ohta et al., 2021). This in turn leads to the recruitment of the γ-tubulin ring complex, which nucleates microtubules (Ohta et al., 2021; Fong et al., 2008; Choi et al., 2010; Dictenberg et al., 1998; Moritz et al., 1995). As the mitotic spindle forms, the centrioles and the PCM become each embedded in the spindle poles, where minus-end binding proteins accumulate (Akhmanova and Steinmetz, 2019).

Centrosomes play an important role in the regulation of the mitotic spindle size (Dudka and Meraldi, 2017). In early *C. elegans* and zebrafish embryos, spindle size directly scales with centrosome size (Greenan et al., 2010; Rathbun et al., 2020). Moreover, previous studies in *Chlamydomonas reinhardtii*, *Ceanorahbditis elegans*, and human cells show that cells with an unequal number of centrioles generate half-spindles of unequal size (Keller et al., 2010; Greenan et al., 2010; Dudka et al., 2019; Tan et al., 2015). In the extreme case, human cells with one centriole in one spindle pole and no centriole in the other spindle pole form highly asymmetric spindles that differ up to 20% in their half-spindle length (Dudka et al., 2019). A half-spindle length asymmetry affects the position of the cytokinetic furrow leading to asymmetric cell daughter size (Dudka et al., 2019; Tan et al., 2015). At the molecular level, spindle size is controlled by several MAPs located at spindle poles (Goshima and Scholey, 2010; Dumont and Mitchison, 2009; Lacroix and Dumont, 2022). These include ch-TOG, CLASP or EB1, which regulate microtubule polymerization, but also microtubule severing enzymes, such as katanin, and microtubule depolymerases of the 8- and 13- kinesin families (Kif18A, Kif2A and MCAK) (Reber et al., 2013; Barr and Gergely, 2008; Goshima et al., 2005; Maiato et al., 2005; McNally et al., 2006; Mayr et al., 2007; Gaetz and Kapoor, 2004). These latter enzymes also play an important role in controlling the size of meiotic spindles in different xenopus species, where elevated levels of katanin or Kif2A have been shown to reduce spindle length in a species-dependent-manner (Loughlin et al., 2011; Miller et al., 2019). In human cells and *Xenopus laevis* extracts, the regulation of the mitotic spindle size also depends on the ability of the Aurora-A/TPX2 complex to regulate microtubule nucleation (Bird and Hyman, 2008; Fu et al., 2015; Helmke and Heald, 2014), while in *C. elegans* embryos, spindle size correlates with the ability to recruit microtubule nucleating material at centrosomes (Greenan et al., 2010). Finally, the asymmetric spindle of the one-cell stage leech *Hellobdella robusta*, relies on the downregulation of the microtubule nucleator γ-tubulin on one spindle pole (Ren and Weisblat, 2006).

A final important aspect of centrosome behavior is centrosome age. Old and young centrosomes are segregated in a stereotypical manner in many stem cell divisions, such as the unicellular budding yeast *Saccharomyces cerevisiae,* murine neuroprogenitors, *Drosophila melanogaster* neuroblasts, or *Drosophila melanogaster* male germline stem cell divisions (Chen and Yamashita, 2021; Yamashita et al., 2007; Januschke et al., 2013; Wang et al., 2009; Pereira et al., 2001). This centrosome-age dependent distribution has been linked to the asymmetric assembly of the primary cilium and its associated hedgehog signaling in the daughter cells in mice (Anderson and Stearns, 2009), and to cellular senescence in yeast (Shcheprova et al., 2008). Centrosome age can also affect the relative half-spindle length in asymmetric stem cell divisions. In *Drosophila melanogaster* neuroblasts, the half-spindle associated to the young centrosome, which is inherited by the stem cell, is longer than the one associated to the old centrosome (Fuse et al., 2003; Roubinet et al., 2017). In somatic human tissue culture cells, we and others have found that centrosome age affects the resolution of polar chromosomes in prometaphase and the segregation of non-disjoint chromosomes in anaphase, due to differential kinetochore-microtubule stability, leading to the preferential inheritance of non-disjoint chromosomes in daughter cells inheriting the old centrosome (Gasic et al., 2015; Colicino et al., 2019). However, whether centrosome age also affects spindle size in somatic tissue culture cells, and thus the relative spindle size symmetry, is not known. Here, we use human epithelial and fibroblastic cell lines to show that centrosome age imposes a subtle but persistent mitotic spindle size asymmetry, and that this spindle size asymmetry translates into an equivalent size asymmetry of the nascent daughter cells. At the mechanistic level, we demonstrate that the polar chromosome asymmetry and the spindle size asymmetry can be experimentally uncoupled, pointing to the existence of multiple spindle asymmetries. Finally, at the molecular level we demonstrate that size asymmetry depends on proteins required for microtubule nucleation that are themselves asymmetrically distributed on the old and young centrosomes. Our study therefore highlights how even in somatic cells thought so far to divide symmetrically, centrosome age breaks spindle and cell division symmetries in multiple ways.

## RESULTS

### Centrosome age breaks spindle size symmetry in cells thought to divide symmetrically

To investigate the contribution of centrosome age to spindle size symmetry we used non-cancerous human retina pigment epithelial cells immortalized with telomerase (hTert-RPE1, called RPE1 hereafter) expressing the centriolar marker GFP-centrin1. In a previous study that did not take centrosome age in account, we had reported that the bulk population of RPE1 cells has on average symmetric spindles (Dudka et al., 2019). Here we analyzed spindle size (a)symmetry versus centrosome age in cells with different centriole configurations: non-treated WT cells containing 2 centrioles at each pole (hereafter called 2:2 cells); cells with one centriole at one spindle pole and no centriole on the other pole as positive control for spindle asymmetry (hereafter caller 1:0 cells); cells without any centrioles as negative control (hereafter called 0:0 cells), and cells with 1 centriole at each spindle pole, since daughter centrioles have also been implicated in spindle length control (hereafter called 1:1 cells; Fig. 1A) (Tan et al., 2015; Dudka et al., 2019). To obtain such centriole configurations, we inhibited centriole duplication with the Plk4 inhibitor centrinone for 24 (1:1), 48 (1:0) or 72 hours (0:0; Fig. 1A) (Wong et al., 2015).

**Figure 1.**
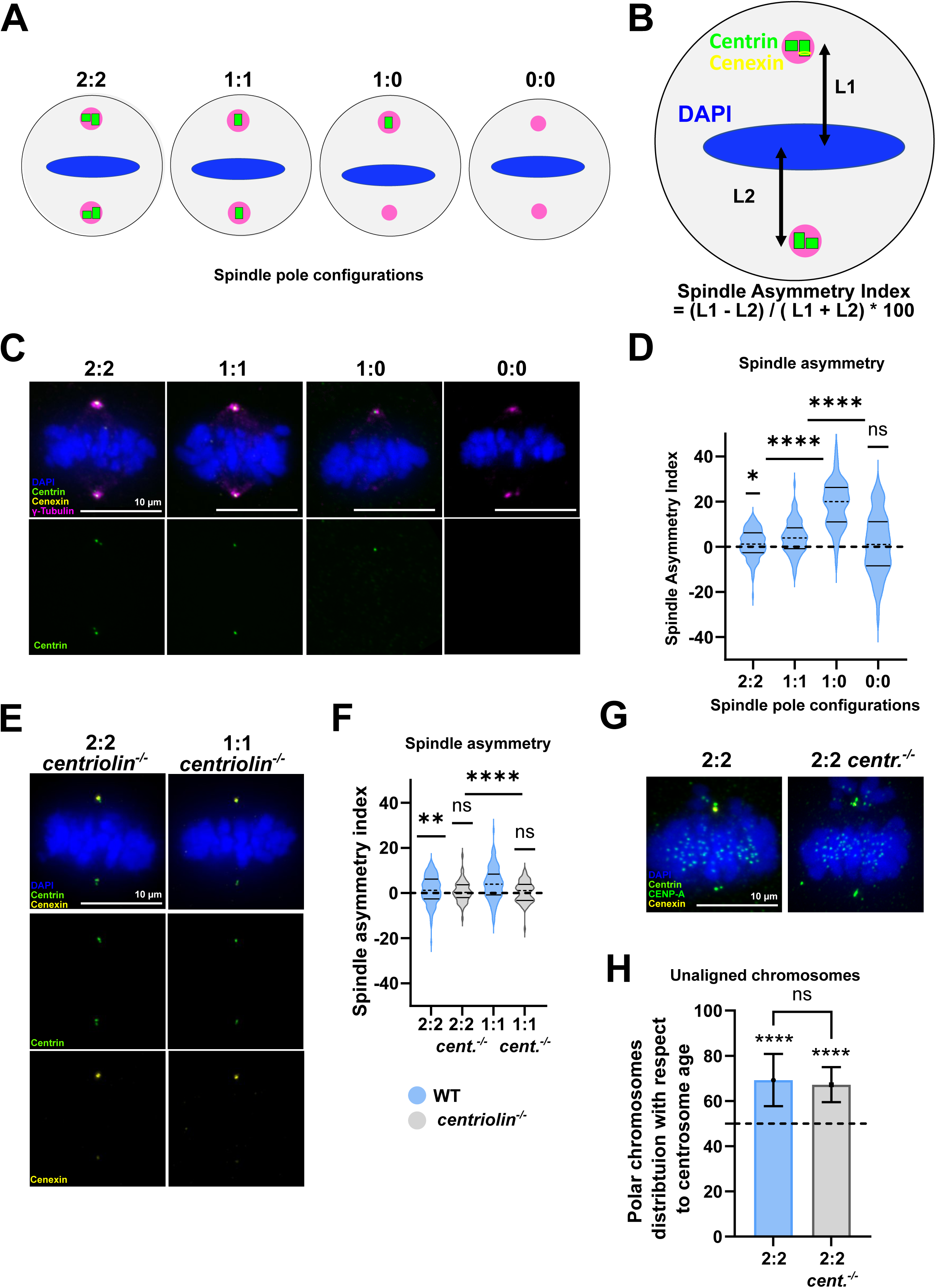
Centrosome age breaks spindle symmetry in RPE1 cells. (**A**) Schematic representation of RPE1 metaphase cells displaying different centriole numbers (green) at spindle poles (magenta). Control non-treated cells contain 2 centrioles at each pole (2:2), cells treated with centrinone during 24h, 48h and 72h display respectively, one centriole at each pole (1:1), one centriole at one pole (1:0) or no centriole (0:0). (**B**) Scheme to explain the calculation of the Spindle Asymmetry Index (SAI) (DAPI). L1 to the distances from the spindle pole with the cenexin-enriched (yellow) old centrosome to the center of the DAPI-stained (blue) metaphase plate (measured in 3D). L2 refers to the distance between the other spindle pole to the metaphase plate. In 0:0 cells, the two spindle poles were assigned randomly. (**C**) Immunofluorescence images of 2:2, 1:1, 1:0 and 0:0 RPE1 GFP-centrin1 metaphase cells. Cells are stained with cenexin (yellow) and γ-tubulin (magenta) antibodies, DAPI (blue) and GFP- centrin1 (green). (**D**) Quantifications of the SAI of 2:2, 1:1, 1:0 and 1:0 cells. The dashed line and full lines of each condition represent the mean and SD respectively. Each condition was compared to a theoretical distribution centered at 0 (dashed line). For 2:2 cells SAI = 1.4% ± 6.2%, n=133 cells, p = 0.0104; for 1:1 cells = 4.1 ± 7.3%, n=96 cells, p <0.0001; for 1:0 cells = 19.3% ± 11.2%, n=111 cells, p <0.0001; and for 0:0 cells = 0.9% ± 13.3%, n=77 cells, p = 0.54 in one-sample t-tests. (**E**) Immunofluorescence images of 2:2 and 1:1 *centriolin^-/-^*RPE1 metaphase cells, stained with DAPI (blue), GFP-centrin1 (green) and cenexin antibody (yellow). (**F**) SAI quantifications of 2:2 and 1:1 (light grey) *centriolin^-/-^* RPE1 cells vs. WT RPE1 (light blue) cells. SAI for 2:2 *centriolin^-/-^* RPE1 cells = 0.7% ± 4.7%, n=56 cells, p = 0.3573; and for 1:*1 centriolin^-/-^* RPE1 cells = 0.4% ± 4.7%, n=65 cells, p = 0.4229 in one-sample t-tests. (**G**) Immunofluorescence images of 2:2 GFP-centrin1 and 2:2 *centriolin^-/-^* RPE1 metaphase cells treated with nocodazole and stained with cenexin (yellow) and CENP-A (green) antibodies, DAPI (blue) and GFP-centrin1 (green). The dashed line represents a symmetrical distribution of the polar chromosomes between the two poles. (**H**) Quantification of the percentage of unaligned chromosomes distributed with respect to centrosome age. Means of 69% ± 12% (from 223 polar chromosomes in 86 cells) 2:2 cells, and 67% ± 8% (from 203 polar chromosomes in 169 cells) 2:2 *centriolin^-/-^* cells. P-values of < 0.0001 for both conditions using binomial tests to compare individual conditions to a random distribution of 50%. P-value of 0.8902 using a Fisher’s exact test to compare the two conditions. All scale bars = 10μm.

Cells were fixed, stained for γ-tubulin to visualize spindle poles, and for the subdistal appendage marker cenexin to differentiate between the old and the young centrosome (Ishikawa et al., 2005). We next measured at the single cell level the two half-spindle lengths (distance from the γ-tubulin signal at spindle pole to the center of the DAPI mass) and plotted the corresponding Spindle (A)symmetry Index (SAI hereafter) versus centrosome age. A positive value indicated a longer half-spindle associated to the old centrosome (Fig. 1B-D). Consistent with previous studies, 1:0 cells assembled highly asymmetric spindles (mean SAI of 19.3% ± 11.2%, n=111 cells, p < 0.0001), while in contrast 0:0 cells displayed a broad range of SAI centered around 0 (mean SAI of 0.9 ± 13.3%, n=77 cells, p = 0.5414; Fig. 1C and D) (Dudka et al., 2019). 2:2 and 1:1 cells had slightly asymmetric SAI distribution with means of 1.4% ± 6.2%, n=133 cells (2:2) and 4.1 ± 7.3%, n=96 cells (1:1) (Fig. 1C and D). A statistical analysis of these distributions confirmed that half-spindles associated to the old centrosomes were statistically significantly longer in both 2:2 and 1:1 cells (p = 0.0104 and p = < 0.0001; Fig. 1C and D).

If centrosome age dictates spindle size asymmetry, we reasoned that it should depend on the subdistal appendages, the main structural difference between the grandmother at old centrosomes and the mother centriole at young centrosomes. We therefore tested whether the abrogation of subdistal appendages rescued the symmetry of the mitotic spindles in 2:2 and 1:1 cells. Since we had to use cenexin as a marker for centrosome age, we could not deplete this most proximal subdistal appendage protein. Instead, we used RPE1 cells in which centriolin, the SDA protein downstream of cenexin, was deleted by CRISPR. This enabled us to still differentiate between old and young centrosome, while removing most subdistal appendage proteins (Mazo et al., 2016). When we measured the SAI in *centriolin^-/-^* cells, we found symmetric spindles in both in 2:2 and 1:1 cells (0.7% ± 4.7%, n=56 cells, p = 0.3573) and 1:1 (0.4% ± 4.7%, n=65 cells, p = 0.4229; Fig. 1E and F). We conclude that even in cells thought to divide symmetrically, centrosome age breaks the symmetry of the mitotic spindle size. Moreover, the fact that 1:1 cells showed a persistently higher spindle size asymmetry raised the possibility that the presence of daughter centrioles dampens centrosome-age dependent size asymmetry.

Since we and others had previously shown that the resolution of polar chromosomes (unaligned chromosomes located behind spindle poles) in prometaphase displays an asymmetry that depends on cenexin (Gasic et al., 2015; Colicino et al., 2019), we next tested whether loss of centriolin and its downstream factors also restores the symmetry of polar chromosomes. Surprisingly, the distribution of polar chromosomes in *centriolin^-/-^* cells remained equally asymmetric as in WT cells (67% ± 11.5% vs 69% ± 11.5% of polar chromosomes associated to the old centrosome, N = 203 and 223 polar chromosomes, n=169 and 86 cells respectively, p = 0.8902; Fig. 1H). This indicates that the centrosome-age dependent asymmetries in terms of polar chromosomes and half-spindle sizes can be uncoupled and that the specific molecular mechanisms governing these processes must differ.

### Centrosome age breaks the symmetry of daughter cell size

In human cells, the position of the metaphase plate influences the position of the actomyosin contractile ring during the cytokinesis (Dudka et al., 2019; Tan et al., 2015). Therefore, an off-center positioning of the metaphase plate of the spindle itself can lead to asymmetric daughter cells sizes (Tan et al., 2015). We therefore probed by immunofluorescence daughter cell size (a)symmetry versus centrosome age in late telophase 2:2, 1:1, and 0:0 cells (Fig 2A). In both 2:2 and 1:1 cells we found that daughter cell size was significantly larger in the cell inheriting the old centrosome (1.9% ± 5.5%, n=66 cells, p = 0.0067; and 4.5% ± 8.2%, n=55 cells, p = 0.0001; Fig. 2B). In contrast, in 0:0 cells daughter cell sizes were symmetric (daughter cell size asymmetry of −0.7% ± 9.5%, n=58 cells, p = 0.6049; Fig. 2B). To exclude that the observed daughter cell size asymmetry originates from an asymmetric position of the spindle itself, we also measured the average distance between old or young centrosome and the cell cortex in metaphase, and found no significant difference (6.8 ± 1.2 μm and 7.1 ± 1.5 μm for the old and young centrosomes respectively, n=44 cells, p = 0.2135, Fig. 2C and D). Finally, to confirm that the asymmetry in daughter cell size depends on centrosome age, we quantified this daughter size asymmetry in 1:1 *centriolin^-/-^*RPE1 cells, and found no asymmetry (−0.2% ± 5.2%, n=53 cells, p = 0.8334; Fig. 2E and F). We conclude that by breaking spindle size symmetry, centrosome age also leads to asymmetric daughter cell sizes.

**Figure 2.**
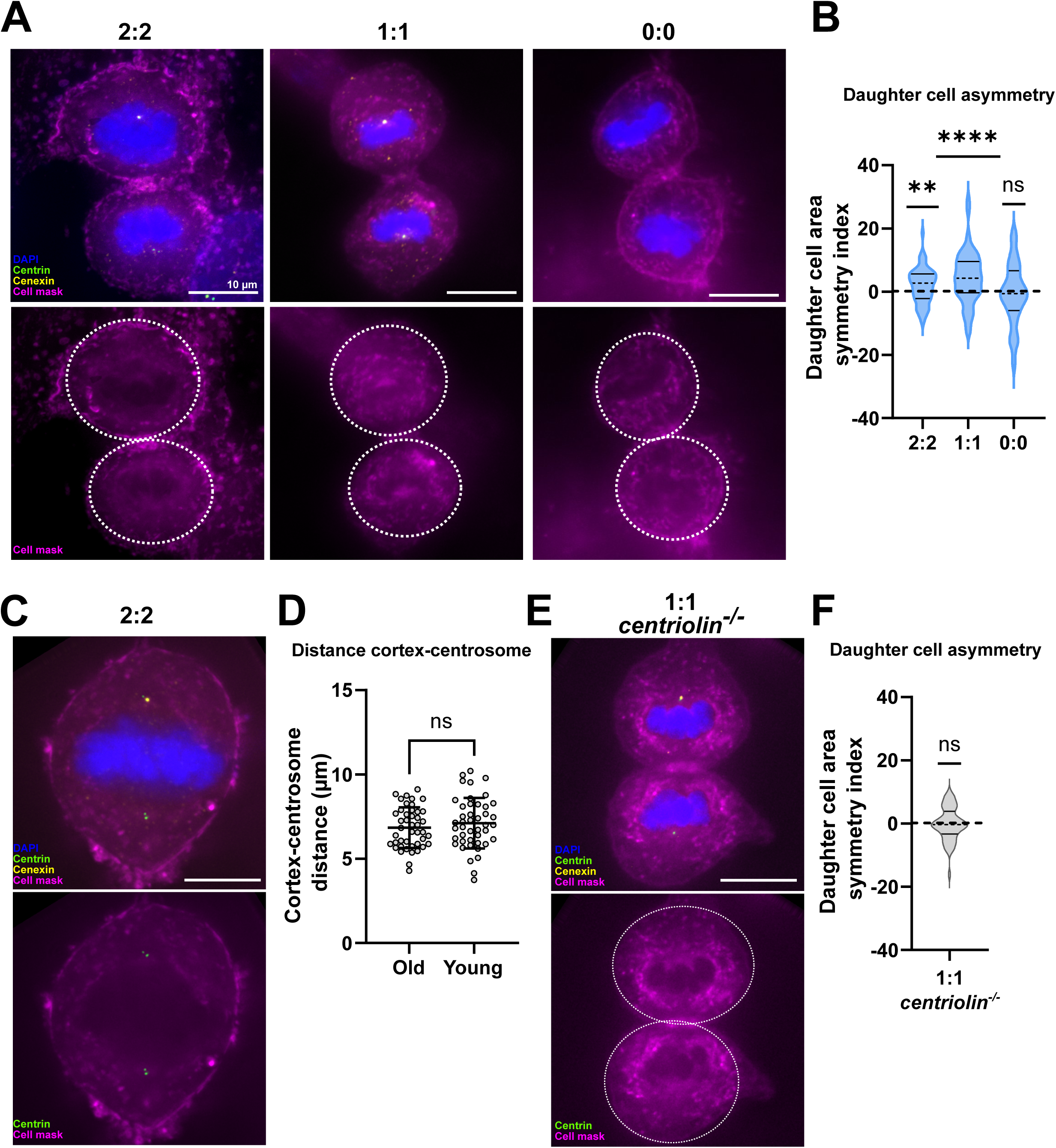
Centrosome-age depedent asymmetric spindles asymmetric daughter cell size. (**A**) Immunofluorescence images of 2:2, 1:1 and 0:0 RPE1 GFP-centrin1 late telophase cells. Cells are stained with DAPI (blue), GFP-centrin1 (green), cenexin (yellow) antibody and the cell mask (magenta). (B) Quantification of the daughter cell area symmetry index. For 2:2 cells = 1.9% ± 5.5%, n=66 cells, p = 0.0067; for 1:1 cells = 4.5 ± 8.2%, n=56 cells, p = 0.0001, and for 0:0 cells =-0,7% ± 9.5%, n=58 cells, p = 0.6049 in one-sample t-tests. (**C**) Immunofluorescence images of 2:2 and 1:1 RPE1 GFP-centrin1 metaphase cells stained with DAPI (blue), GFP-centrin1 (green), cenexin antibody (yellow) and the cell mask (magenta). (**D**) Quantification of the cortex-centrosome distances. Distance means of 6.8 ± 1.2 μm and 7.1 ± 1.5 μm for the old and young centrosomes respectively (44 cells, p = 0.2135, ns in paired t-test. (**E**) Immunofluorescence images of a 1:1 *centriolin^-/-^* RPE1 late telophase cell stained with DAPI (blue), GFP-centrin1 (green), cenexin antibody (yellow) and the cell mask (magenta). (**F**) Quantification of the daughter cell area symmetry index of 1:1 *centriolin^-/-^* RPE1 cells = −0.2% ± 5.2%, n=53, p = 0.8334, in one-sample t-test. All scale bars = 10μm.

### Centrosome age breaks spindle size symmetry via microtubule nucleation

We next aimed to identify the molecular mechanisms by which centrosome age breaks spindle size symmetry and generally test how centrosomes affect spindle size. We first quantified the relative abundance of a series of centrosome and spindle pole proteins that have been implicated in spindle size control. For each cell we plotted, based on 3D immunofluorescence images, the relative protein distribution asymmetry between the two poles versus the spindle asymmetry index (Fig. 3A-B, and Supplementary Fig. 1). We carried out this analysis on 2:2 and 1:1 cells, which show a centrosome-age dependent spindle asymmetry, but also in 1:0 and 0:0 cells. This allowed us to investigate whether the protein abundance distribution correlates specifically with centrosome-age dependent spindle asymmetry, or more generally to spindles with an asymmetric distribution of centrosomes, or no centrosomes at all. From these plots we extracted two key values: the Pearson’s R correlation coefficient, which reflects how reliably the protein abundance distribution correlates with spindle (a)symmetry, and the slope of the linear regression, which indicates the potential degree by which a protein might contribute to spindle (a)symmetry (Fig. 3B, Supplementary Fig. 1 and Table 1). We considered proteins as potential drivers of spindle (a)symmetry when the correlation coefficient was statistically significant (p < 0.01), and the slope of the linear regression exceeded 0.15.

**Figure 3.**
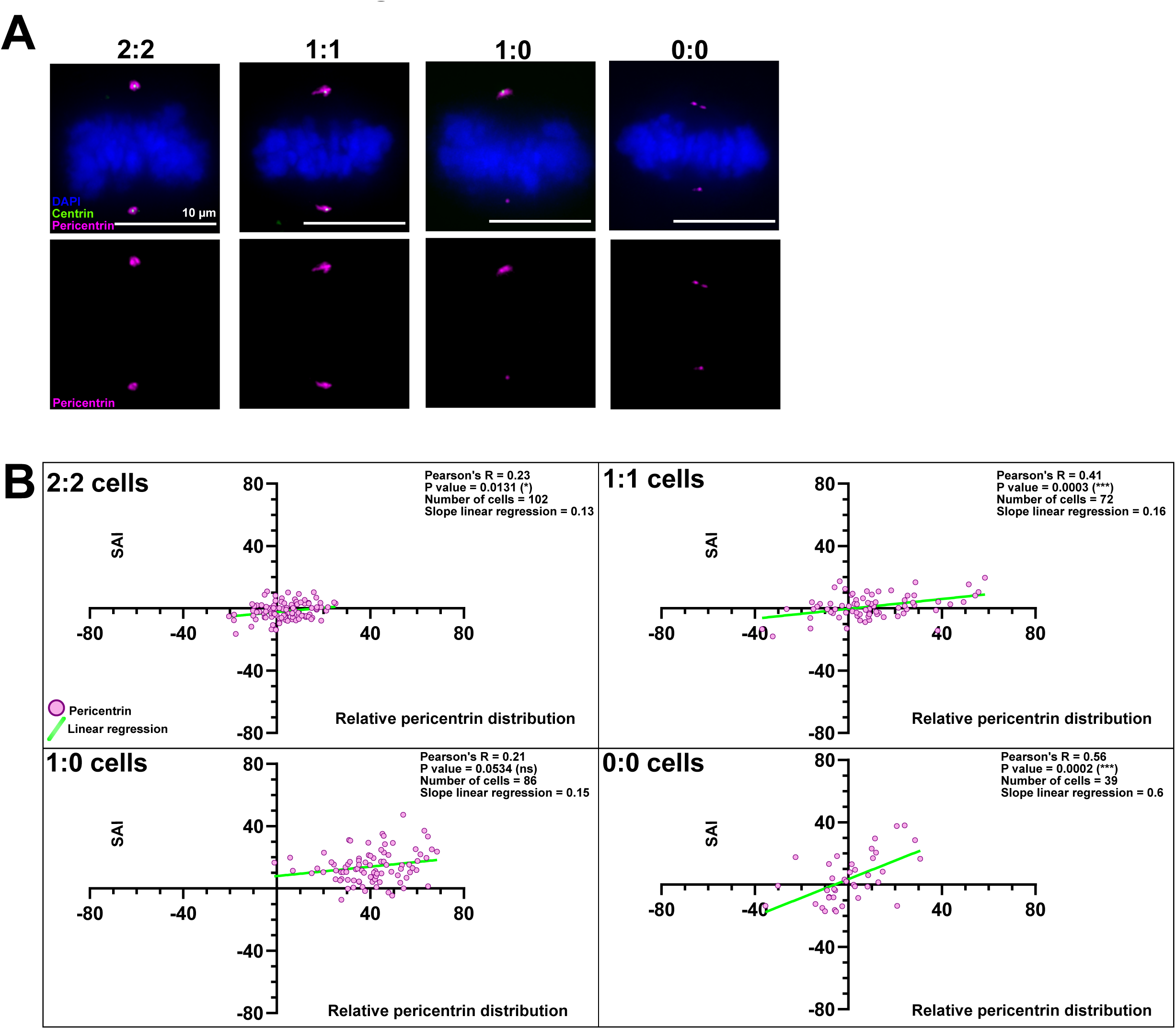
The spindle pole abundance of proteins regulating microtubule nucleation scales with half-spindle size. (**A**) Immunofluorescence images of 2:2, 1:1, 1:0 and 0:0 RPE1 GFP-centrin1 metaphase cells stained with DAPI (blue), GFP-centrin1 (green), and pericentrin antibody (magenta). Scale bars = 10 μm. (**B**) Correlation plots between the relative (B) pericentrin distribution (X axis) and the SAI (Y axis) for 2:2, 1:1, 1:0 and 0:0 cells. Dots represent single cell values. The light green line indicates the slope of the linear regression. For each plot the Pearson correlation coefficient, its associated P-value, the number of cells analyzed and the slope of the linear regression are indicated. The data for all the proteins can be found in the table 1.

**Table 1.**
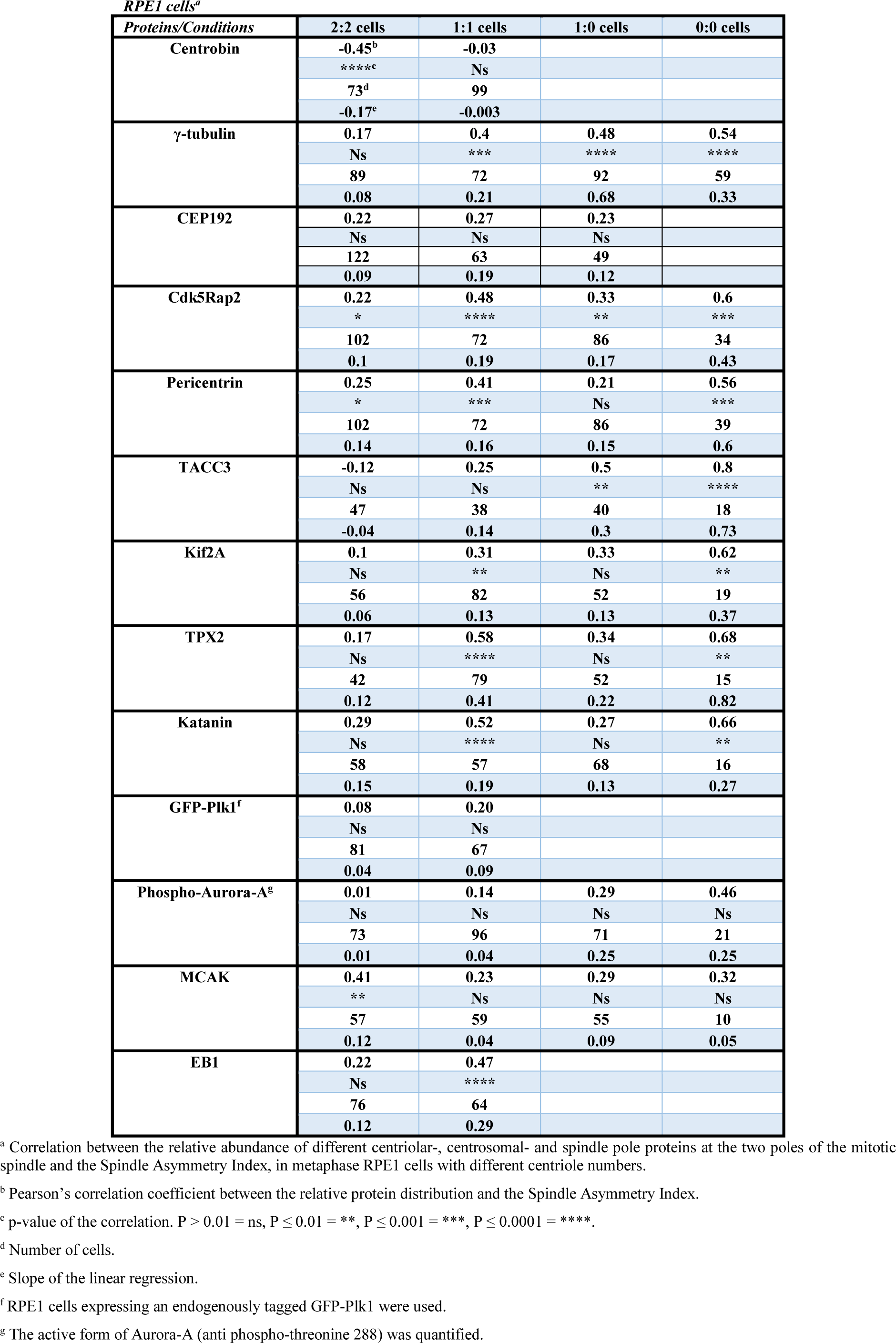
Identification of potential drivers of spindle (a)symmetry in RPE1 cells.

| *Proteins/Conditions* | **2:2 cells** | **1:1 cells** | **1:0 cells** | **0:0 cells** |
| --- | --- | --- | --- | --- |
| **Centrobin** | **-0.45<sup>b</sup>** | **-0.03** |  |  |
|  | ****<sup>c</sup> | Ns |  |  |
|  | **73<sup>d</sup>** | **99** |  |  |
|  | **-0.17<sup>e</sup>** | **-0.003** |  |  |
| **γ-tubulin** | **0.17** | **0.4** | **0.48** | **0.54** |
|  | Ns | *** | **** | **** |
|  | **89** | **72** | **92** | **59** |
|  | **0.08** | **0.21** | **0.68** | **0.33** |
| **CEP192** | **0.22** | **0.27** | **0.23** |  |
|  | Ns | Ns | Ns |  |
|  | **122** | **63** | **49** |  |
|  | **0.09** | **0.19** | **0.12** |  |
| **Cdk5Rap2** | **0.22** | **0.48** | **0.33** | **0.6** |
|  | * | **** | ** | *** |
|  | **102** | **72** | **86** | **34** |
|  | **0.1** | **0.19** | **0.17** | **0.43** |
| **Pericentrin** | **0.25** | **0.41** | **0.21** | **0.56** |
|  | * | *** | Ns | *** |
|  | **102** | **72** | **86** | **39** |
|  | **0.14** | **0.16** | **0.15** | **0.6** |
| **TACC3** | **-0.12** | **0.25** | **0.5** | **0.8** |
|  | Ns | Ns | ** | **** |
|  | **47** | **38** | **40** | **18** |
|  | **-0.04** | **0.14** | **0.3** | **0.73** |
| **Kif2A** | **0.1** | **0.31** | **0.33** | **0.62** |
|  | Ns | ** | Ns | ** |
|  | **56** | **82** | **52** | **19** |
|  | **0.06** | **0.13** | **0.13** | **0.37** |
| **TPX2** | **0.17** | **0.58** | **0.34** | **0.68** |
|  | Ns | **** | Ns | ** |
|  | **42** | **79** | **52** | **15** |
|  | **0.12** | **0.41** | **0.22** | **0.82** |
| **Katanin** | **0.29** | **0.52** | **0.27** | **0.66** |
|  | Ns | **** | Ns | ** |
|  | **58** | **57** | **68** | **16** |
|  | **0.15** | **0.19** | **0.13** | **0.27** |
| **GFP-Plk1<sup>f</sup>** | **0.08** | **0.20** |  |  |
|  | Ns | Ns |  |  |
|  | **81** | **67** |  |  |
|  | **0.04** | **0.09** |  |  |
| **Phospho-Aurora-A<sup>g</sup>** | **0.01** | **0.14** | **0.29** | **0.46** |
|  | Ns | Ns | Ns | Ns |
|  | **73** | **96** | **71** | **21** |
|  | **0.01** | **0.04** | **0.25** | **0.25** |
| **MCAK** | **0.41** | **0.23** | **0.29** | **0.32** |
|  | ** | Ns | Ns | Ns |
|  | **57** | **59** | **55** | **10** |
|  | **0.12** | **0.04** | **0.09** | **0.05** |
| **EB1** | **0.22** | **0.47** |  |  |
|  | Ns | **** |  |  |
|  | **76** | **64** |  |  |
|  | **0.12** | **0.29** |  |  |

<sup>a</sup> Correlation between the relative abundance of different centriolar-, centrosomal- and spindle pole proteins at the two poles of the mitotic spindle and the Spindle Asymmetry Index, in metaphase RPE1 cells with different centriole numbers.
<sup>b</sup> Pearson's correlation coefficient between the relative protein distribution and the Spindle Asymmetry Index.
<sup>c</sup> p-value of the correlation. P > 0.01 = ns, P ≤ 0.01 = \*\*, P ≤ 0.001 = \*\*\*, P ≤ 0.0001 = \*\*\*\*.
<sup>d</sup> Number of cells.
<sup>e</sup> Slope of the linear regression.
<sup>f</sup> RPE1 cells expressing an endogenously tagged GFP-Plk1 were used.
<sup>g</sup> The active form of Aurora-A (anti phospho-threonine 288) was quantified.

**Table 2.**
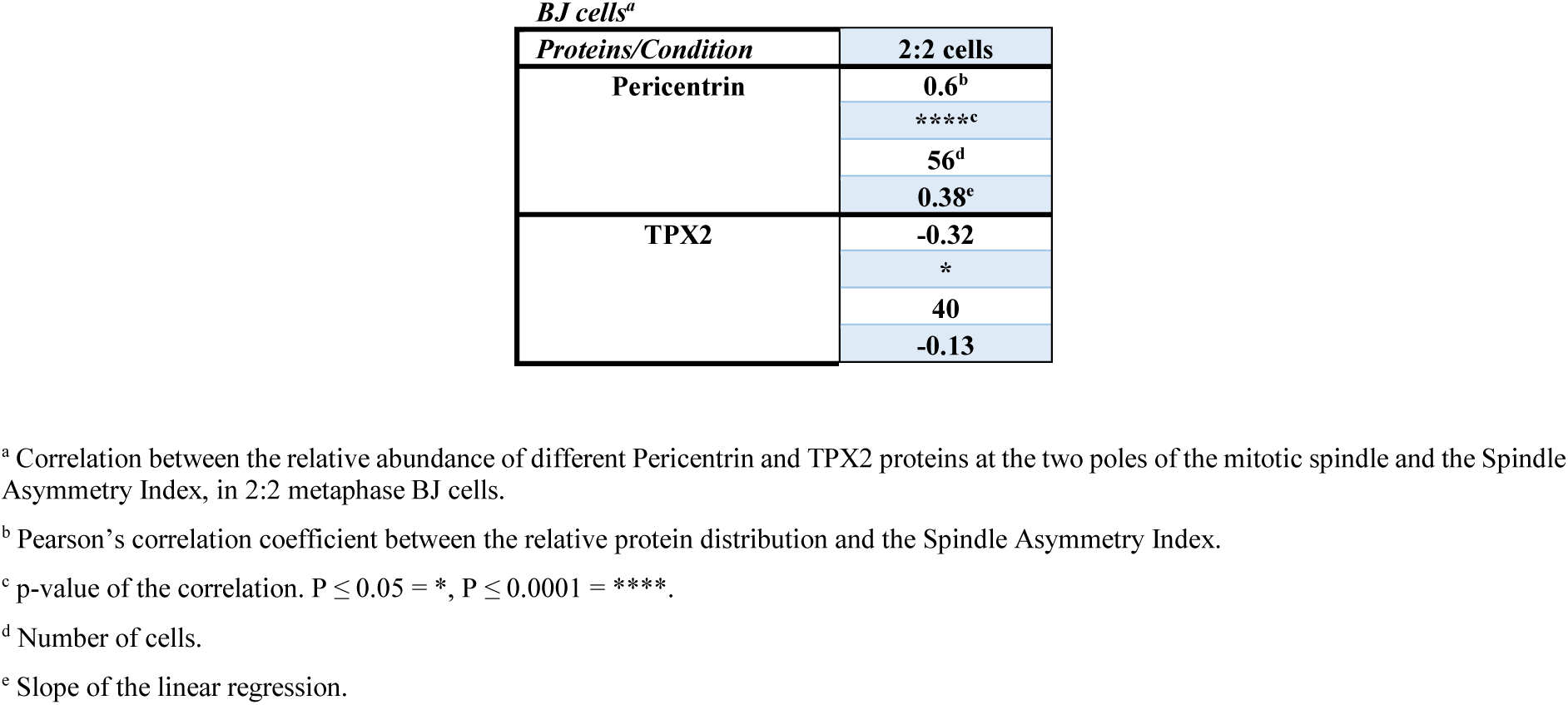
Identification of potential drivers of spindle (a)symmetry in BJ cells.

The list of investigated proteins included centrobin, a daughter centriole-specific protein (Zou et al., 2005; Jeffery et al., 2010); the mitotic kinase Plk1 and Aurora-A (specifically the activated phosphorylated form of Aurora-A) and their centrosomal activator CEP192, which have been implicated in the regulation of spindle microtubule dynamics (Joukov et al., 2014; Barr and Gergely, 2007; Asteriti et al., 2015); pericentrosomal proteins required for microtubule nucleation such as pericentrin, Cdk5Rap2 and γ-tubulin that have been implicated in spindle size regulation (Fong et al., 2008; Choi et al., 2010; Greenan et al., 2010; Ren and Weisblat, 2006; Watanabe et al., 2020); and microtubule dynamics regulators at spindle poles such as the microtubule depolymerases Kif2A and MCAK (Ganem and Compton, 2004; Jang et al., 2009; Domnitz et al., 2012), the microtubule-severing enzyme Katanin (Loughlin et al., 2011; Huang et al., 2021; Guerreiro et al., 2021), as well as the microtubule-associated proteins TPX2 (Bird and Hyman, 2008; Sobajima et al., 2023), TACC3 (Cassimeris and Morabito, 2004; Gergely et al., 2003) and EB1 (Dema et al., 2022).

Amongst the hits we identified in this analysis were pericentrin, Cdk5Rap2 and γ-tubulin, whose protein abundance distribution significantly correlated with the SAI in 1:1, 1:0 and 0:0 cells (Fig. 3B, Supplementary Fig. 1 and Table 1), meaning that at the single cell level the longer half-spindle also tended to display more of those proteins at the spindle pole. This suggested that pericentriolar proteins implicated in microtubule nucleation, might control spindle (a)symmetry both in the presence or absence of centrosomes. Even in 2:2 cells, there was a visible correlation between the relative pericentrin and Cdk5Rap2 distribution and the spindle asymmetry (Table 1). We next tested whether their distribution on the two spindle poles follows centrosome age and whether these proteins contribute to spindle (a)symmetry. We focused our attention in particular on 1:1 cells, which display a sharper spindle asymmetry than 2:2 cells. We found that all three proteins were enriched on the spindle pole associated to the old centrosome (means of 14% ± 13.8%, 13.3% ± 15.7%, and 4.4% ± 6,2% respectively for pericentrin, Cdk5Rap2 and γ-tubulin), and that their depletion via siRNAs abolished the asymmetry of 1:1 cells (SAI means of 0.3% ± 6.1%, −0,3% ± 7%, and 0.9% ± 9.5% respectively for pericentrin, Cdk5Rap2 and γ-tubulin; Fig. 4A-C; validation of all siRNA treatments in Supplementary Fig. 2). This indicated that their presence is necessary to break spindle symmetry in a centrosome age dependent manner. In contrast when we depleted as a negative control centrobin, a protein not known to control spindle size, we found that the spindles were still asymmetric (SAI mean of 3.4% ± 7.74%, n=23 cells, p = 0.0493, in *siCtrl* vs. 4.5% ± 6.1%, n=33 cells, p = 0.0002, in *sicentrobin*; Fig. 4D and E; Supplementary Fig. 2). This indicated that the effects of pericentrin, Cdk5Rap and γ-tubulin on age-dependent spindle symmetry were specific. In contrast, depletion of pericentrin or Cdk5Rap2 did not affect spindle asymmetry in 1:0 cells (γ-tubulin could not be tested, as its depletion disrupted spindle formation in 1:0 cells; Fig. 4F-H). Those findings implied that proteins implicated in microtubule nucleation at centrosomes are specifically required for centrosome-age dependent spindle asymmetry. Consistent with this hypothesis, we found in a microtubule re-nucleation assay that the capacity to nucleate microtubules at centrosomes correlated with their age. Indeed, when microtubules were depolymerized by a 1-hour cold treatment, both centrosomes contained the same quantity of residual microtubules; in contrast, when microtubules were allowed re-nucleate for 15s in warm medium, more tubulin could be found at the old centrosome (Fig. 4I and J). Finally, when we tested whether the loss of centriolin cells affected the distribution of these proteins, we found that γ-tubulin, but not pericentrin, lost its asymmetric distribution in *centriolin^-/^*^-^ (Fig. 4K; Cdk5Rap2 levels could not be tested due to antibody incompatibilities). We conclude that microtubule nucleation at centrosomes depends on centrosome age and contributes to centrosome-age dependent spindle asymmetry.

**Figure 4.**
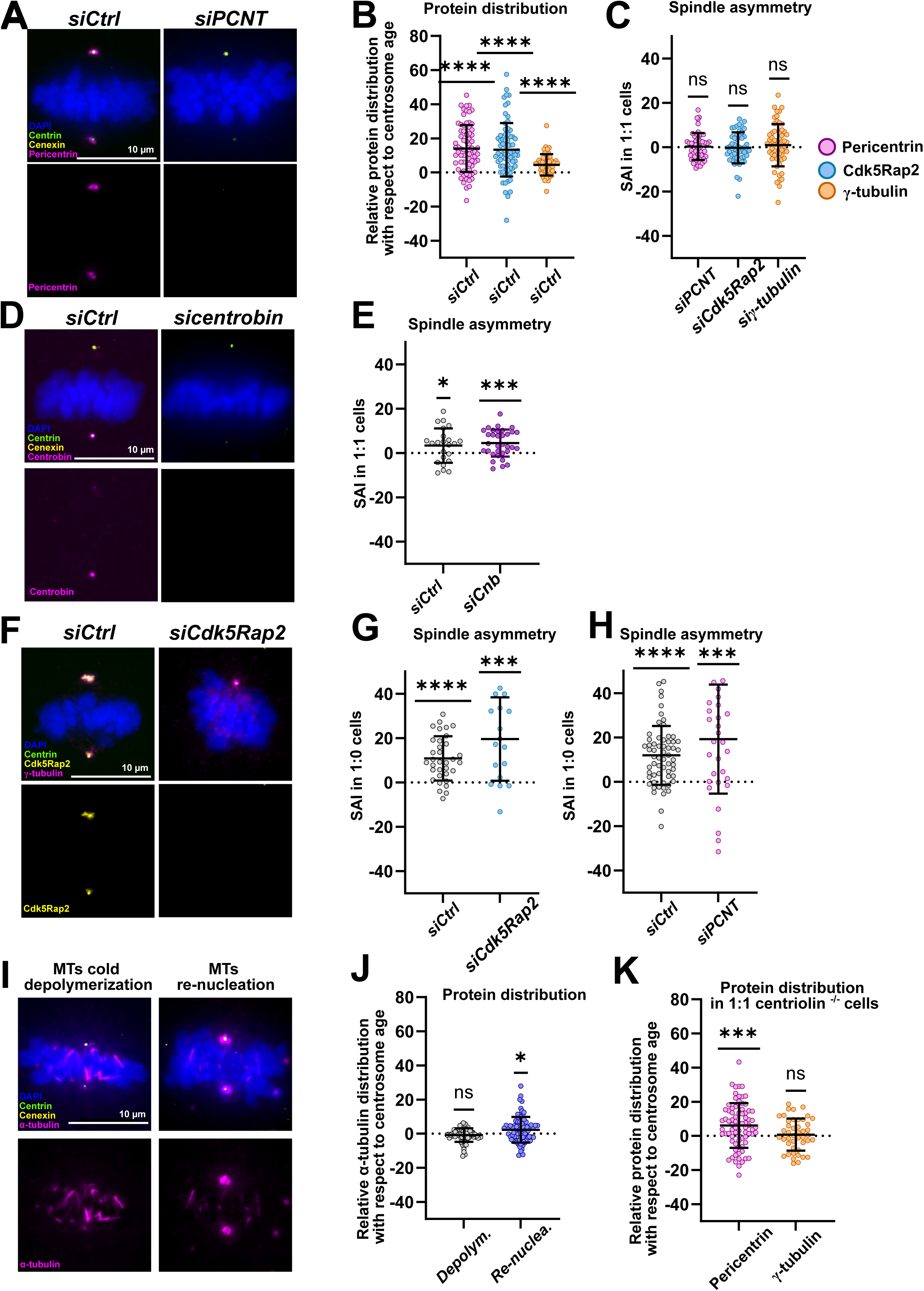
Proteins regulating microtubule nucleation break spindle symmetry. (**A**) Immunofluorescence images of *siCtrl* and *siPCNT*-treated RPE1 GFP-centrin1 1:1 cells stained with DAPI (blue), centrin1-GFP (green), cenexin (yellow) and pericentrin (magenta) antibodies. (**B**) Quantifications of the relative pericentrin, Cdk5Rap2 and γ-tubulin asymmetry index vs. centrosome age (using cenexin as old centrosome marker) in control-depleted 1:1 RPE1 GFP-centrin1 cells. Means of 13.99% ± 13.76%, 13.30% ± 15.67%, and 4.43% ± 6,23% for pericentrin, Cdk5Rap2 and γ-tubulin respectively. P <0.0001 in one-sample t-tests for pericentrin and Cdk5Rap2, and one-sample Wilcoxon test for γ-tubulin. (**C**) Quantifications of the SAI after *siPCNT*, *siCdk5Rap2* and *siγ-tubulin* treatment in 1:1 RPE1 GFP-centrin1 cells. Means of 0.27% ± 6.06%, −0.29% ± 6.96%, and 0.92% ± 9.52% for pericentrin, Cdk5Rap2 and γ-tubulin depletion respectively. P-values of respectively 0.8246, 0.8902 and 0.3277 for *siPCNT*, *siCdk5Rap2* and *siγ-tubulin* in one-sample t-tests tests. (**D**) Immunofluorescence images of *siCtrl* and *sicentrobin*-treated 1:1 RPE1 GFP-centrin1 cells stained with DAPI (blue), GFP-centrin1 (green), cenexin (yellow) and centrobin (magenta) antibodies. (**E**) Quantifications of the SAI in *siCtrl and sicentrobin*-treated in RPE1 GFP-centrin1 cells. SAI means 3.4% ± 7.74%, n=23 cells, p = 0.0493, in *siCtrl* cells and 4.5% ± 6.1%, n=33 cells, p = 0.0002, in *sicentrobin* cells. One-sample t-tests were used for statistical analyses. (**F**) Immunofluorescence images of *siCtrl* and *siCdk5Rap2*-treated 1:0 RPE1 GFP-centrin1 cells, stained with DAPI (blue), GFP-centrin1 (green), Cdk5Rap2 (yellow) and γ-tubulin (magenta) antibodies. (**G** and **H**) Quantifications of the SAI of (G) *siCtrl and siCdk5Rap2- or (H) siCtrl and siPCNT*-treated 1:0 RPE1 GFP-centrin1 cells. SAI means of (G) 10.8% ± 10%, n= 24 cells, p < 0.0001 in *siCtrl* cells and 19.6% ± 18.8%, n= 18 cells, p = 0.0007, in *siCdk5Rap2* cells. One-sample Wilcoxon tests were used for statistical analyses. SAI means of (H) 11.9% ± 13.2%, n= 61 cells, p < 0.0001, in *siCtrl* cells and 19.2% ± 24.6%, n= 32 cells, p < 0.0001 in *siCdk5Rap2* cells. One-sample t-tests were used for statistical analyses. (**I**) Immunofluorescence images of 1:1 RPE1 GFP-centrin1 cells after 1h of cold-depolymerization (left) and 15s after re-nucleation (right). Cells are stained with DAPI (blue), GFP-centrin1 (green), cenexin (yellow) and α-tubulin (magenta) antibodies. (**J**) Quantifications of the relative α-tubulin distribution at centrosomes vs. centrosome age. After the cold depolymerization, mean of −0.8% ± 4%, n=75 cells, p = 0.2044, and after the re-nucleation, mean of 2.3% ± 7.5%, n=76 cells, p = 0.0256. One-sample Wilcoxon tests were used for statistical analyses. (**K**) Quantifications of the relative protein distribution of pericentrin and γ-tubulin with respect to centrosome age in 1:1 *centriolin^-/-^* RPE1 cells. Means of 6.1% ± 13.05% and 0.73% ± 9.42% for pericentrin, and γ-tubulin respectively. P-values > 0.0001 and 0.6059 for pericentrin and γ-tubulin respectively in one-sample t-tests. All Scale bars = 10μm.

### Centrosome age breaks spindle size symmetry via a pericentrin-TPX2 axis

From our analysis we identified a second group of potential hits, which included the microtubule depolymerase Kif2A, the microtubule severing enzyme katanin and the regulators of microtubule dynamics TPX2 and TACC3. While Kif2A, katanin and TPX2 were typically enriched on the spindle pole of the longer half-spindle in 1:1, 1:0 and 0:0 cells, TACC3 only showed such a correlation in 1:0 and 0:0 cells (Table 1). This suggested that TACC3 may only regulate spindle length/symmetry in the context of centrosome-free spindle poles and may not depend on centrosome age. This observation is consistent with its localization at spindle poles in human oocytes, as a component of the centrosome-free oocyte microtubule organizing centers (Wu et al., 2022).

The higher abundance of a microtubule-depolymerase (Kif2A) and microtubule severing enzyme (katanin) on the longer half-spindle appeared at first counter-intuitive, as one would expect such enzymes to reduce half-spindle length. Nevertheless, this profile could be explained by the known ability of TPX2 to form complexes with Kif2A/CLASP1 and with katanin/WDR62 on the mitotic spindle (Fu et al., 2015; Huang et al., 2021). We therefore reasoned that TPX2, which regulates spindle size on its own (Bird and Hyman, 2008; Fu et al., 2015; Helmke and Heald, 2014; Sobajima et al., 2023), could be a critical factor regulating spindle asymmetry. To test this hypothesis, we quantified the SAI in both 1:1 and 1:0 cells after depleting TPX2 by siRNA (validation see Supplementary Fig. 3A; note that to visualize the spindle poles in *siTPX2* cells we used antibodies against the minus-end binding protein NuMa). Our measurements indicated a rescue of spindle symmetry in 1:1 cells compared to a control depletion (1.7% ± 5.6%, n=55 cells, p = 0.0389, in *siControl* cells and 1.3% ± 9.3%, n=79 cells, p = 0.2342, ns, in *siTPX2* cells) and a significant decrease of the spindle asymmetry in 1:0 cells (Fig. 5A-D). These results imply that TPX2 in contrast to the pericentrin/Cdk5Rap2/g- tubulin module contributes to the spindle asymmetry in both 1:1 and 1:0 cells. Consistent with a role in centrosome-age dependent spindle asymmetry, TPX2 itself localized in an asymmetric manner in wild-type RPE1 2:2 cells, but not in *centriolin^-/-^* RPE1 2:2 cells, indicating that its localization is influenced by centrosome age (means of 3% ± 6.5%, n=36 cells, p = 0.0210, in WT cells vs. 0.7% ± 5%, n=55 cells, p = 0.5917 in *centriolin^-/-^* cells; Fig. 5E).

**Figure 5.**
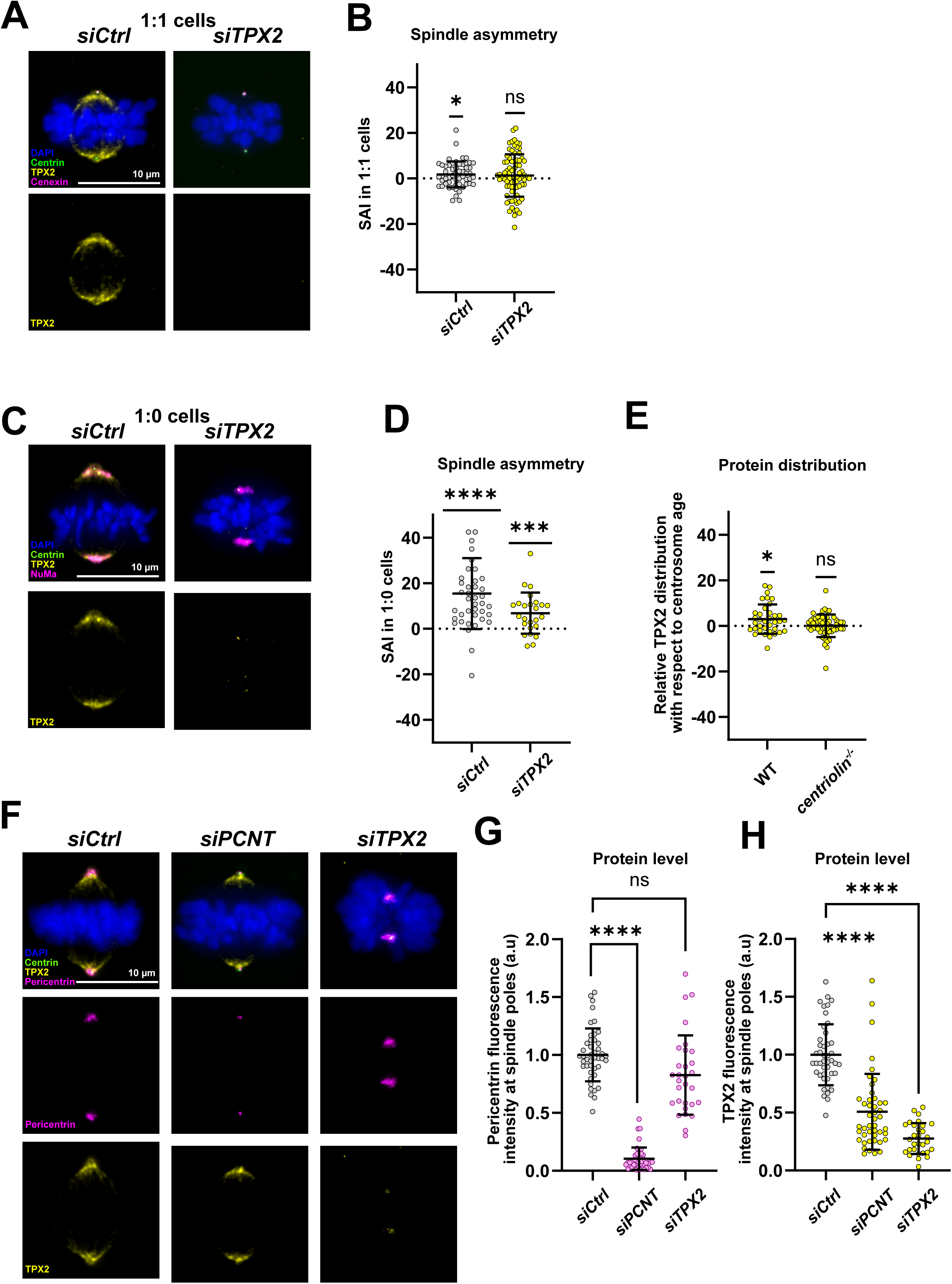
A pericentrin-TPX2 axis drives the formation of asymmetric spindles. (**A**) Immunofluorescence images of *siCtrl* and *siTPX2*-treated 1:1 RPE1 GFP-centrin1 cells stained with DAPI (blue), GFP-centrin1 (green), cenexin (yellow) and TPX2 (magenta) antibodies. (**B**) Quantifications of the SAI of *siCtrl and siTPX2*-treated 1:1 RPE1 GFP-centrin1 cells. SAI means of 1.7 % ± 5.6%, n=55 cells, p = 0.0389, in *siCtrl* cells and 1.3% ± 9.3%, n=79 cells, p = 0.2342, in *siTPX2* cells. One-sample t-tests were used for statistical analyses. (**C**) Immunofluorescence images of *siCtrl* and *siTPX2-*treated 1:0 RPE1 Centrin1-GFP cells, stained with DAPI (blue), GFP-centrin1 (green), cenexin (yellow) and TPX2 (magenta) antibodies. (**D**) Quantifications of the SAI of *siCtrl and siTPX2-* treated 1:0 RPE1 Centrin1-GFP cells. SAI means of 15.4% ± 15.6%, n=41 cells, p < 0.0001, in *siCtrl* cells and 6.8% ± 9%, n=25 cells, p = 0.0009, in *siTPX2* cells. One-sample t-tests were used for statistical analyses. (**E**) Quantification of the TPX2 asymmetry vs centrosome age in 2:2 RPE1 GFP-centrin1 WT and *centriolin^-/-^* cells. Relative TPX2 distribution means of 2.9% ± 6.5%, n=36, p = 0.0210, in WT cells and 0.1% ± 5%, n=55, p = 0.5917, in *centriolin^-/-^* cells. (**F**) Immunofluorescence images of *siCtrl, siPCNT and siTPX2-*treated 2:2 RPE1 GFP-centrin1 cells stained with DAPI (blue), GFP-centrin1 (green), TPX2 (yellow) and pericentrin (magenta) antibodies. (**G** and **H**) Quantification of (G) pericentrin and (H) TPX2 fluorescence intensity at spindle poles in *siCtrl (n=44 cells)*, *siPCNT (n=45 cells)* and *siTPX2-*treated (n=30 cells) 2:2 RPE1 GFP-centrin1 cells. p < 0.0001, ****, in one-way Kruskal-Wallis with Dunnett’s multiple comparisons tests. All scale bars = 10μm.

Given that TPX2 and the pericentriolar proteins pericentrin/Cdk5Rap2/γ-tubulin are both required for centrosome age dependent spindle asymmetry we used siRNA depletions to test whether they affect each other’s recruitment at spindle poles (Fig. 5F), as to our knowledge no interaction between TPX2 and pericentrin or Cdk5Rap2 has been reported so far. Quantitative immunofluorescence indicated that pericentrin depletion reduced TPX2 intensity at spindle poles by 50%, while TPX2 depletion did not significantly change pericentrin abundance (Fig. 5G and H). We conclude that pericentrin contributes to TPX2 recruitment at spindle poles and that it acts upstream of TPX2 in the centrosome-age dependent breaking of spindle symmetry.

### Pericentrin is also present on daughter centrioles

One of the surprising results was the finding that 1:1 cells lacking daughter centrioles displayed a stronger spindle asymmetry than wild-type 2:2 RPE1 cells. This implied that the presence of daughter centrioles dampens the symmetry breaking imposed by centrosome age. Daughter centrioles are generally considered to be immature; however, our and others work have shown that in conditions where daughter centrioles prematurely disengage during mitosis, such daughter centrioles can organize spindle poles and recruit γ-tubulin (Wilhelm et al., 2019; Logarinho et al., 2012). We therefore used a high-resolution Airyscan microscope to visualize whether daughter centrioles can also recruit the pericentriolar seed protein pericentrin under wild-type conditions. Consistent with previous high-resolution studies (Mennella et al., 2012; Lawo et al., 2012), we found that pericentrin formed rings around the mature grandmother and mother centrioles. Nevertheless, the same images also revealed incomplete pericentrin rings around both daughter centrioles in metaphase cells, indicating that they indeed can recruit pericentriolar material, and may thus dampen the spindle asymmetry imposed by the grandmother centriole (Fig. 6A).

**Figure 6.**
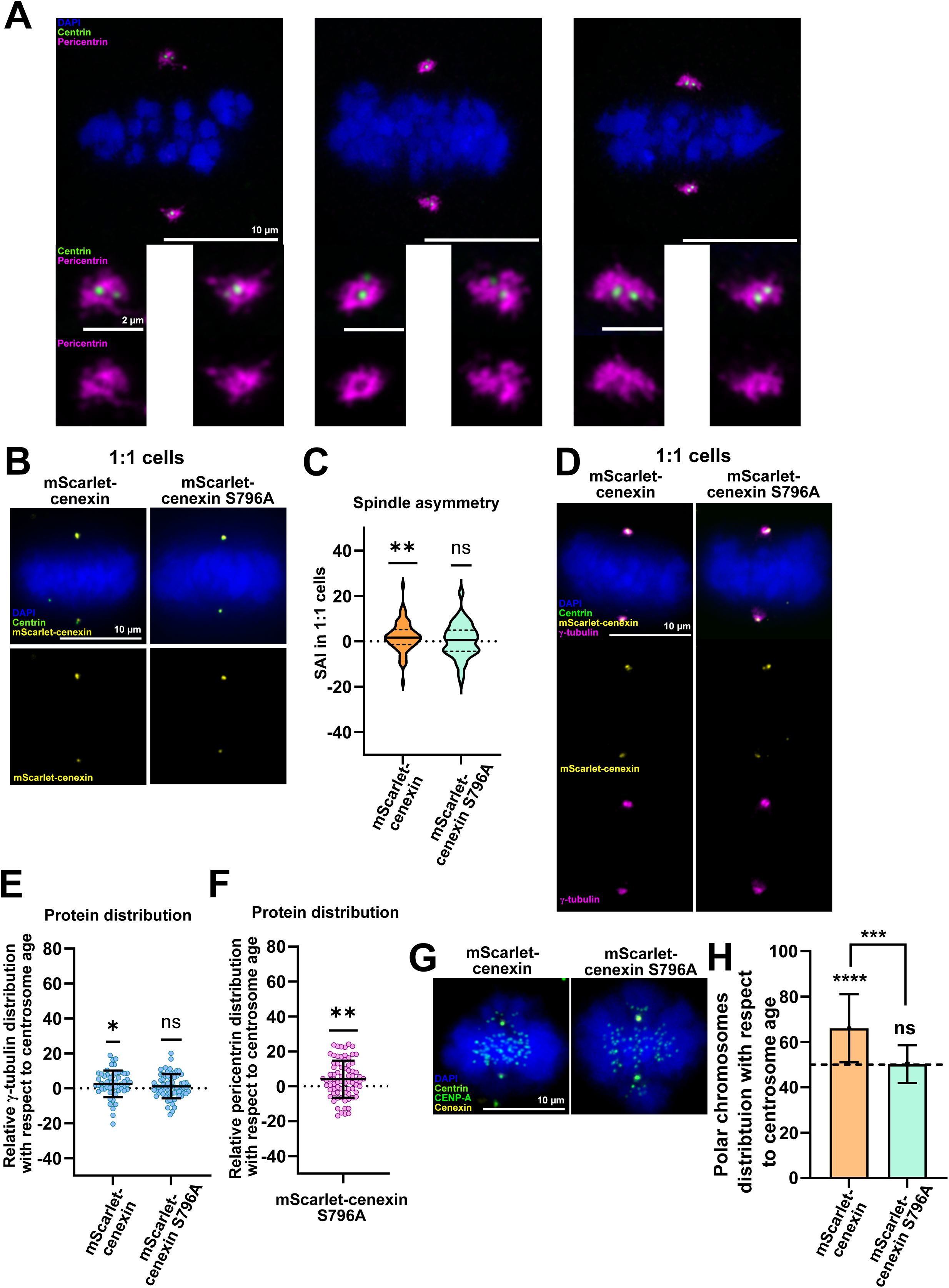
The phosphorylation of cenexin S796 drives the formation of asymmetric spindles. (**A**) High-resolution immunofluorescence images of metaphase RPE1 GFP-centrin1 cells, stained with DAPI (blue), GFP-centrin1 (green), pericentrin (magenta) antibody. Scale bars = 10μ. Insets show a maximum intensity projection of pericentrin at centrosomes. Scale bars = 2μ. (**B**) Immunofluorescence images of 1:1 metaphase RPE1 GFP-centrin1 cells expressing either WT mScarlet-cenexin or S796 mScarlet-cenexin, stained with DAPI, GFP-centrin1 (green), and mScarlet-cenexin antibody (yellow). Scale bar = 10μ. (**C**) Quantification of the SAI in 1:1 RPE1 GFP-centrin1 cells expressing either WT mScarlet-cenexin or S796 mScarlet-cenexin. WT-cenexin SAI mean: 2.0% ± 6.5%, n=73 cells, p = 0.0044 in one-sample Wilcoxon test compared to a symmetric null hypothesis. S796A cenexin SAI mean: 0.35± 7.5%, n= 75 cells, p = 0.7096 in one-sample t-test. (**D**) Immunofluorescence images 1:1 metaphase RPE1 GFP-centrin1 cells expressing either WT mScarlet-cenexin or S796 mScarlet-cenexin, stained with DAPI, GFP-centrin1 (green), mScarlet-cenexin (yellow) and γ-tubulin (magenta) antibodies. Scale bar = 10μ. (**E**) Quantifications of the relative γ-tubulin distribution with respect to centrosome age in 1:1 RPE GFP-centrin1cells expressing either WT or S796A mScarlet-cenexin. In WT-cenexin cells relative distribution mean of 2.6 ± 7.6%, n= 40 cells, p = 0.020 and in S796A cenexin cells relative distribution mean of 1.2 ± 6.1%, n= 55 cells, p = 0.149 in one-sample Wilcoxon test. (**F**) Quantifications of the relative pericentrin distribution with respect to centrosome age in 1:1 RPE GFP- centrin1 mScarlet-cenexin S796A cells. Relative distribution mean of 4.1± 10.6%, n= 75 cells, p = 0.0014 in one-sample t-test. (**G**) Immunofluorescence images of 1:1 RPE GFP-centrin1 cells expressing either WT or S796A mScarlet-cenexin metaphase cells treated with nocodazole and stained with cenexin (yellow) and CENP-A (green) antibodies, DAPI (blue) and GFP-centrin1 (green). (**H**) Quantification of the percentage of unaligned chromosomes distributed with respect to centrosome age. Means of 69% ± 15% (from 267 polar chromosomes in 96 cells) for mScarlet-cenexin cells, and 50% ± 8% (from 235 polar chromosomes in 75 cells) for mScarlet-cenexin S796A cells. P-values of < 0.0001 and 1 for mScalert-cenexin and mScarlet-cenexin S796A cells respectively using binomial tests to compare individual conditions to a random distribution. P-value of 0.0003 (***) using a Fisher’s exact test to compare the two conditions.

### Cenexin-bound Plk1 controls both spindle size and polar chromosome asymmetries

A second puzzling result was the fact that pericentrin, Cdk5Rap2 and γ-tubulin showed asymmetric distribution, being enriched on the older centrosome, but that the main regulator of pericentriolar recruitment, the mitotic kinase Plk1 (Lee and Rhee, 2011; Lane and Nigg, 1996; Conduit et al., 2014), did not correlate with spindle size asymmetry (Table 1). We reasoned, however, that a subpool of Plk1 is recruited specifically on the grandmother centriole once the C-terminus part of cenexin is phosphorylated by CDK1 on S796 (Soung et al., 2009). We therefore tested whether the abrogation of this phosphorylation site would abrogate spindle size asymmetry. We introduced in RPE1 *cenexin^-/-^*cells either WT cenexin tagged with mScarlet (mScarlet-cenexin) or a mutant in which the Plk1 binding site has been mutated (S796A mScarlet-cenexin). Both cenexin versions localized predominantly to one centrosome (presumably the older centrosome), and none of the two mutants affected mitotic timing (defined as the time between nuclear envelope breakdown and anaphase onset, Figure 6B and Supplementary Figure 3). Their effect on spindle size symmetry, however, was very different. While WT mScarlet-cenexin 1:1 displayed an asymmetric half-spindle lengths (2% ± 6.5%, n=73 cells, p = 0.0044; Fig. 6C), 1:1 cells expressing mScarlet-cenexin S796A harbored symmetric spindles (0.4% ± 7.5%, n=75 cells, p = 0.7096; Fig. 6C). Consistently, γ-tubulin distribution was slightly asymmetric in 1:1 cells expressing WT mScarlet-cenexin (2.4± 7.6%, n= 40 cells, p = 0.020), but symmetric in S796A mScarlet-cenexin 1:1 cells (1.2± 6.1%, n= 55 cells, p = 0.149; Figure 6D and E). Finally, we asked whether the absence of the Plk1 binding-site on cenexin also affects polar chromosome asymmetry, as one would predict based on previous studies showing that this asymmetry depends both on cenexin and Plk1 (Gasic et al., 2015; Colicino et al., 2019). Our quantification indicated that 1:1 cells expressing WT mScarlet-cenexin displayed an asymmetric distribution of polar chromosomes; in contrast, cells expressing S796A mScarlet-cenexin displayed a symmetric distribution of polar chromosomes (69% ± 15% vs 50% ± 8% of polar chromosomes associated to the old centrosome for mScarlet-cenexin and mScarlet-cenexinS796A respectively, N = 267 and 235 polar chromosomes, n=96 and 75 cells, p = 0.8902; Fig. 6G and H). We conclude that the cenexin-bound pool of Plk1 breaks spindle symmetry both in terms of half-spindle sizes and in terms of polar chromosome distribution.

### Centrosome age also breaks spindle size symmetry in human fibroblast cells

To investigate if this spindle size asymmetry is RPE1 cell-dependent, or a more general feature of tissue human cells, we carried out equivalent experiments with the fibroblastic, non-transformed human cell line BJ-hTERT (called BJ cells hereafter). We first quantified the SAI of the spindles of wild-type metaphase BJ cells (2:2 cells), using cenexin as a marker for centrosome age, and found that BJ cells also displayed asymmetric half-spindle lengths, with longer half-spindles associated to the old centrosome (3.1% ± 7.6%, n=106 cells, p < 0.0001; Fig. 7A-B). This spindle size asymmetry correlated with an asymmetric daughter cell size in 1:1 cells, as we found that the daughter cells inheriting the grandmother centriole was statistically larger than the one inheriting the mother centriole (5.7% ± 9.2%, n=36 cells, p = 0.0007; Fig. 7C-D).

**Figure 7.**
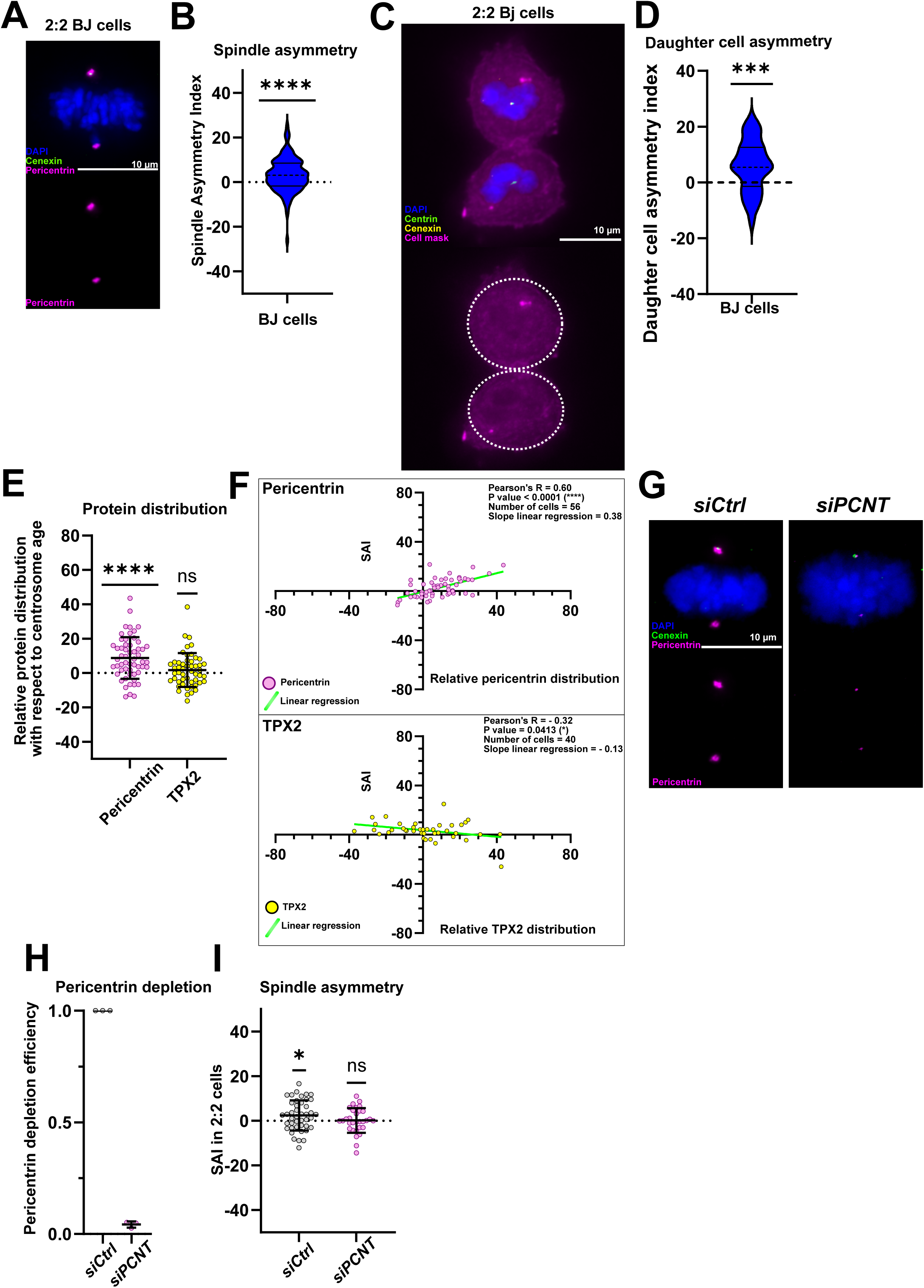
Centrosome age also breaks spindle symmetry in human fibroblast cells. (**A**) Immunofluorescence image of 2:2 BJ metaphase cell stained with DAPI (blue), cenexin (green), and pericentrin (magenta) antibodies. (**B**) Quantification of the SAI in 2:2 BJ cells. SAI mean of 3.1% ± 7.6%, n=106 cells, p < 0.0001 in one-sample t-test. (**C**) Immunofluorescence images of 1:1 BJ late telophase cell stained with DAPI (blue), centrin (green), cenexin (yellow) antibodies and the cell mask (magenta). (**D**) Quantification of the daughter cell area asymmetry of 1:1 BJ cells. Daughter cell size asymmetry of 5.7% ± 9.2%, n=36 cells, p = 0.0007 in one-sample t-test. (**E**) Quantifications of the relative protein distribution of pericentrin and TPX2 with respect to centrosome age in 2:2 BJ cells. Relative pericentrin distribution of 8.7% ± 12.1%, n=56 cells, p < 0.0001. Relative TPX2 distribution of 1.7% ± 9.9%, n=51 cells, p = 0.4329. A one-sample t-test was used for pericentrin statistical analysis and a one sample Wilcoxon-test was used for TPX2 statistical analysis. (**F**) Correlation between the relative pericentrin/TPX2 distribution (X axis) and the SAI (Y axis) in 2:2 BJ cells. Each dot represents a cell for which the relative pericentrin/TPX2 distribution and the SAI were measured. The Pearson correlation coefficient and its associated P-value, the number of cells analyzed as well as the slope of the linear regression (light green line) are indicated for each condition. (**G**) Immunofluorescence images of *siCtrl-* and *siPCNT-*treated 2:2 BJ cells, stained with DAPI (blue), cenexin (green), and pericentrin (magenta) antibodies. (**H**) Quantification of pericentrin depletion efficiency of *siCtrl-* and *siPCNT-* treated 2:2 BJ cells. Pericentrin fluorescence intensity means of 100%, n=33 cells in *siCtrl* and 4.2% ± 1.4%, n=30 in *siPCNT* cells. (**I**) Quantifications of the SAI of *siCtrl- and siPCNT-*treated 2:2 BJ cells. SAI means of 2.4% ± 6.8%, n= 43 cells, p = 0.0250, *, in *siCtrl* cells and 0.1% ± 5.6%, n= 43 cells, p = 0.8988 in one-sample t-tests. All scale bars = 10μm.

We next investigated the distribution of the two main regulators identified in RPE1 cells, pericentrin and TPX2. Quantitative immunofluorescence using cenexin as a marker for centrosome age indicated that on average pericentrin was asymmetrically distributed, whilst TPX2 displayed a symmetric distribution (8.7% ± 12.1%, n=56 cells, p < 0.0001; and 1.7% ± 9.9%, n=51 cells, p = 0.22, respectively for pericentrin and TPX2; Fig. 7E). At the single cell level, the abundance of pericentrin was highly correlated with the SAI (Pearson correlation coefficient of 0.60, slope value of 0.38), whilst the abundance of TPX2 did not correlate with spindle asymmetry (Figure 7F). Consistent with our data showing an important role of pericentriolar material, pericentrin depletion led to symmetric spindles in 2:2 BJ cells (2.4% ± 6.8%, n=43 cells, p = 0.0250 for *siCtrl*, and 0.1% ± 5.6%, n=31 cells, p = 0.8988 for *siPCNT*; Figure 7G-H). We conclude that a centrosome-age linked spindle symmetry is a general feature of human cell lines, present in epithelial and fibroblastic cells and that this spindle asymmetry in both cases depends on the pericentriolar material.

## DISCUSSION

It has long been thought that human generic tissue culture cells divide symmetrically with half-spindles of equal lengths. Here, we demonstrate that centrosome age breaks spindle size symmetry even in cells thought to divide symmetrically, and that this results in the formation of two daughter cells of unequal size (see model figure 8). Our data further indicate that this symmetry breaking is related but different to the previously identified polar chromosome asymmetry. At the molecular level we show that in RPE1 cells centrosome age-dependent spindle symmetry depends on the unequal recruitment of proteins involved in the control of microtubule nucleation, via an axis involving cenexin-bound Plk1, the subdistal appendages, the recruitment of pericentriolar material, and TPX2 (Figure 8). Finally, our data point to an unexpected role of daughter centrioles in dampening spindle asymmetry.

**Figure 8:**
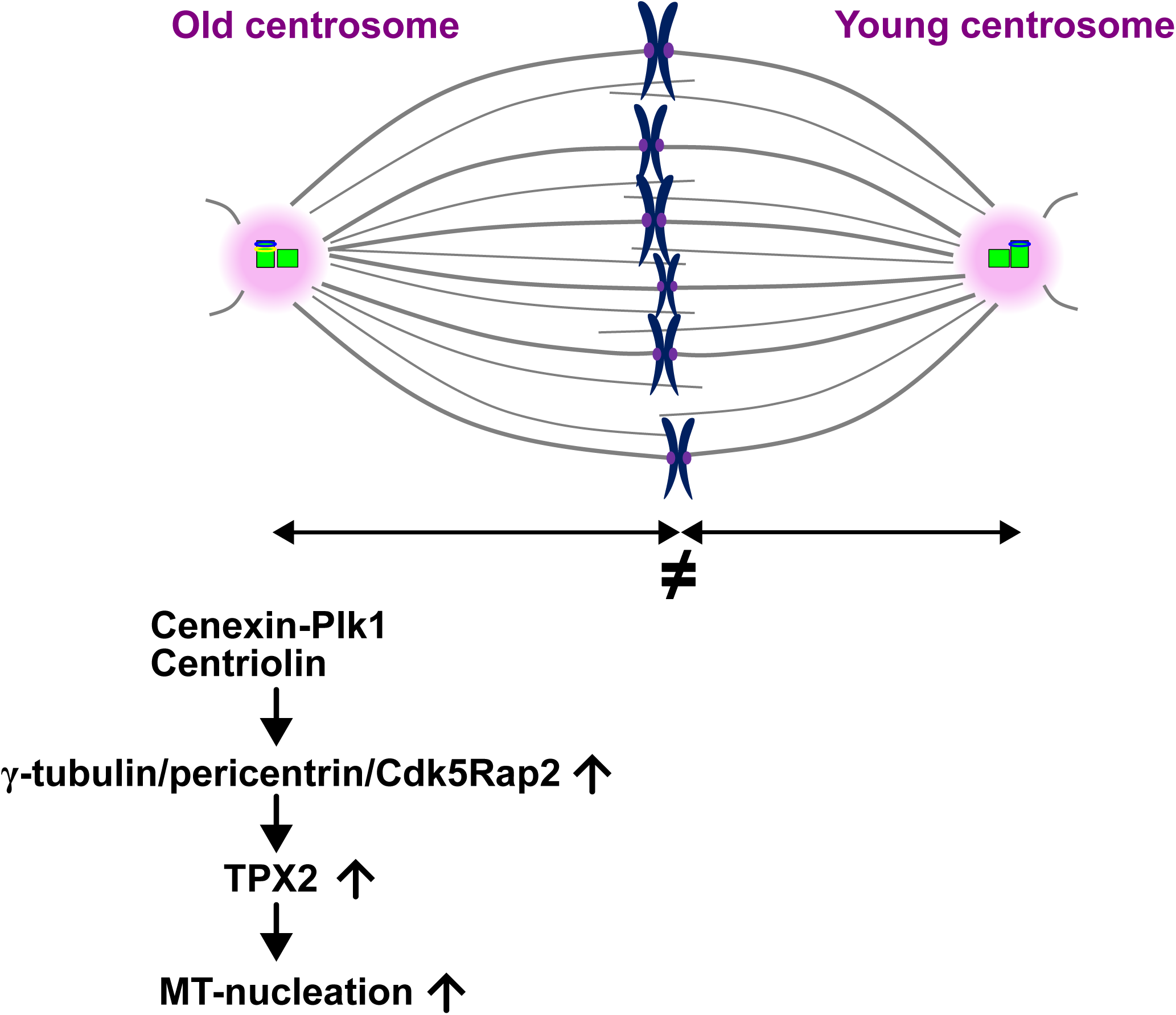
Schematic model for the control of spindle asymmetry in RPE1 cells. We postulate that spindle size asymmetry is controlled in a centrosome age dependent manner via the cenexin-bound pool of Plk1 and centriolin, which both regulate the abundance of pericentriolar material required for microtubule nucleation, which itself regulates the abundance of TPX2.

It is known since a long time that old and young centrosomes display different behaviors in stem cell divisions (Chen and Yamashita, 2021; Yamashita et al., 2007; Januschke et al., 2013; Wang et al., 2009; Pereira et al., 2001), and that they can affect the relative half-spindle lengths in Drosophila neuroblasts (Fuse et al., 2003; Roubinet et al., 2017). Here, we show that centrosome age also breaks the symmetry of half-spindle lengths in cells thought to divide symmetrically, creating a bias towards longer half-spindles associated to the old centrosome. These results differ from our previous measurements in HeLa (Tan et al., 2015) and RPE1 cells (Dudka et al., 2019), which pointed to symmetric spindles. One important difference is that these studies relied on the brightest GFP-centrin1 signal as a marker for the old centrosome, which we found to only partially coincide with the cenexin-positive, old centrosome in RPE1 cells (see supplementary Figure 5). We therefore hypothesize that this inconsistency masked the subtle half-spindle length asymmetry found in this study. Nevertheless, this asymmetry is found in two non-transformed human cells of epithelial and fibroblastic origin, arguing for a general phenomenon amongst human cells. It also biases the placement of the cell division plane, resulting in asymmetric daughter cell sizes. Even though the difference is subtle, we note that previous studies found that such differences in cell size can impact the length of the ensuing G1 phase and the probability of cell death (Kiyomitsu and Cheeseman, 2013).

With regard to previous studies in somatic tissue culture cells, our data show that the half-spindle size asymmetry is related, yet different to the previously identified polar chromosome asymmetry (Gasic et al., 2015; Colicino et al., 2019). Both depend on the cenexin-bound pool of Plk1, but only the spindle size asymmetry depends on centriolin at sub-distal appendages. This indicates that both types of asymmetries are independent of each other, as they can be uncoupled, and point to, at least in part, differing molecular mechanism imposing these asymmetries. Whether the centrosome-age dependent spindle size asymmetry has a direct function in generic cell divisions, or whether it is a passive consequence of a generally higher microtubule nucleation capacity at old centrosomes, remains unclear. The study of “symmetric” cells expressing only S796A cenexin and comparing them to isogenic “asymmetric” cells expressing WT cenexin, will therefore be important in future experiments aiming to study the potential long-term consequences of centrosome-age dependent asymmetries.

Our correlation analysis of protein abundances and spindle asymmetry index at the single cell level allowed to identify potential drivers of spindle asymmetry, that could then be (in)validated by siRNA treatments. These included proteins that had been previously found to display asymmetries, such such as pericentrin, Cdk5Rap2 or centrobin (Tan et al., 2015; Gasic et al., 2015; Colicino et al., 2019; Roux-Bourdieu et al., 2022), but also other proteins such as γ-tubulin or TACC3. Our functional data indicate that proteins implicated in microtubule nucleation are key for the centrosome-age dependent spindle size asymmetry. Specifically, our data shows that centrosome-age spindle asymmetry depends on proteins regulating microtubule nucleation (Moritz et al., 1995; Bird and Hyman, 2008; Dictenberg et al., 1998; Petry et al., 2013; Fong et al., 2008; Choi et al., 2010), that are themselves asymmetrically distributed (TPX2/pericentrin/Cdk5Rap2/γ-tubulin). Our functional data indicate that microtubule nucleation at centrosomes is asymmetric. These data are consistent with previous studies in *Ceanorhabditis elegans*, showing that spindle size scales with the size of the microtubule-nucleating pericentriolar material (Greenan et al., 2010), the fact that the leech *Hellobdella robusta* downregulates γ-tubulin on one spindle pole to create an asymmetric cell division (Ren and Weisblat, 2006), and the fact that asymmetric stem cell divisions in *Drosophila melanogaster* are characterized by a strong difference in microtubule nucleation capacity between the old and young centrosome (Yamashita et al., 2007; Januschke et al., 2013). Our data also suggest that the exact molecular mechanisms that creates an asymmetric microtubule nucleation might vary depending on the context, as we find that TPX2 does not correlate with spindle asymmetry in BJ cells. In contrast in the context of the highly asymmetric 1:0 spindles, TPX2 depletion has a much stronger effect than pericentrin depletion. Our results also point to the existence of other, centrosome-independent mechanisms that may control spindle size symmetry, such as TACC3, whose abundance we find to only correlate with spindle size asymmetry in spindles with centrosome-free spindle poles. Such mechanisms might be particularly important in centrosome-free systems, such as meiotic oocytes. Consistent with this hypothesis, we note that TACC3 and TPX2, whose abundance also strongly correlated with spindle size asymmetry in 1:0 cells, are particularly important for spindle assembly in murine and human oocytes (Brunet et al., 2008; Wu et al., 2022).

At the detailed molecular level, we find that centrosome age affects microtubule nucleation via several pathway. First, we find that spindle asymmetry and asymmetric recruitment of γ-tubulin depends on the cenexin-bound subpool of Plk1, consistent with data showing that this phosphorylation site contributes to the efficient recruitment of pericentrin and γ-tubulin (Soung et al., 2009). Second using *centriolin^-/-^*cells, we show that subdistal appendages in general are necessary for these asymmetries. Since cenexin is still enriched on the old centrosome in *centriolin^-/-^* cells (Mazo et al., 2016), these two pathways must work in parallel. We also show that in RPE1 cells TPX2 recruitment to spindle poles is diminished by half in the absence of pericentrin. TPX2 forms a complex with Aurora-A and the minus-end binding protein NuMA at spindle poles (Kufer et al., 2002; Polverino et al., 2021), but another subpool of TPX2 might interact with the pericentriolar material and might thus be recruited in a differential manner to both spindle poles.

Finally, the fact that 1:1 cells display in all conditions a stronger asymmetry than wild-type 2:2 cells, indicates that daughter centrioles are not just immature passengers of the spindle poles during mitosis, but may have an active role. Such as an active role is consistent with our observation that daughter centrioles can recruit pericentriolar material in mitosis (this study), and the fact that under conditions of premature centriole dis-engagement, daughter centrioles can nucleate and organize spindle microtubules (Wilhelm et al., 2019; Logarinho et al., 2012). Although very speculative at this stage, the fact that we identify an activity that in wild-type cells suppresses centrosome-age spindle asymmetry, could suggest that cells have selected mechanisms that can actively counteract this asymmetry, and thus modulate it according to the context of the different types of cell division.

## MATERIAL AND METHODS

### Cell culture, cell lines and drug treatments

hTert-RPE1, hTert-RPE1 GFP-Plk1 (kind gift of M. Gotta, University of Geneva), hTert-RPE1 GFP-centrin1 (kind gift of A. Khodjakov, New York State Department of Health, Wadsworth Center), hTert-RPE1 *cenexin^-/-^*, hTert-RPE1 *centriolin^-/-^* (both king gift from M.F. Tsou, National Yang-Ming University), and hTert-RPE1 *cenexin^-/-^* GFP-centrin1 Scarlet-Cenexin (this study) and hTert-RPE1 *cenexin^-/-^* GFP-centrin1 Scarlet-Cenexin S796A (this study) cells were cultured in high glucose DMEM (Thermofisher; 41965), supplemented with 10% FCS (LabForce; S1810), and 1% penicillin/streptomycin (Thermofisher; 15140). hTert-BJ (kind gift from R. Medema, Netherlands Cancer Institute) cells were cultured in DMEM:F12 (Thermofisher; 11320033), and supplemented with 10% FCS and 1% penicillin/streptomycin. For live-cell imaging, cells were cultured in eight-well µ-Slide ibidi chambers (Ibidi; 80806) in Leibovitz’s L-15 medium without Phenol Red (Thermofisher; 21083) supplemented with 10% FCS. To visualize DNA, 1 nM of SiR-DNA (Spirochrome) was added in the L-15 medium 4 hours prior imaging. Centrinone (Tocris, UK) (Wang et al., 2015) was added to the culture medium at 300 nM during either 24h, 48h or 72h to obtain 1:1, 1:0 and 0:0 cells respectively. Centrinone containing medium was exchanged twice a day with fresh centrinone aliquots to prevent Plk4 reactivation. To increase the number of polar chromosomes, cells were treated during 2 hours with a low dose of nocodazole (10 ng/ml) (Vasquez et al., 1997, Gasic et al., 2015).

### Lentivirus production and cells transduction

To generate the hTert-RPE1 *cenexin^-/-^* expressing GFP-centrin1 and Scarlet-cenexin, hTert-RPE1 *cenexin^-/-^* GFP-centrin1 cells were transduced with lentivirus containing a plasmid coding mScarlet- cenexin. 4.5×10^6^ 293T cells (ATCC; CRL-11268) were seeded in a 10-cm dish, and transiently transfected for 24h with 10μg psPax2 packaging plasmid (plasmid #12260, Addgene), 5μg pMD2.G envelope plasmid (plasmid #12259, Addgene) and with 15 μg mScarlet-cenexin plasmid (Twist BioScience), in a CaCl2 250 mM and Hebs1X (Hepes 6 g.l^-1^, NaCL 8 g.l^-1^, Na2HPO4 0.1g.l^-1^, dextrose 1 g.l^-1^) medium. After 72h, viruses were harvested from the supernatant and filtered with a 0.4 µm filter. 1×10^6^ hTert-RPE1 *cenexin^-/-^* GFP-centrin-1 cells were transduced with 50 μL virus harvest in a 10-cm dish. After 24h, the virus-containing medium was exchanged with fresh DMEM. Individual positive clones expressing mScarlet-cenexin were sorted by FACS after 72h.

### Site-directed mutagenesis of the cenexin cDNA

To obtain the mScarlet-cenexin S796A construct, we replaced by PCR the UCU codon coding for Serine 796 by an Alanine GCU codon using the mScarlet-cenexin plasmid as template. Site directed mutagenesis was performed according to manufacturer’s instructions (QuickChange II, Agilent, 200523-12). Following the PCR reaction, the template plasmid was digested with *Dpn I*, and we used 10 µL of the reaction mix to transform 50 µL of *E. coli* Mach1 (Invitrogen, C8620-03) by heat shock. The sequence was verified by sequencing. Virus production and cell transduction were performed as described for the mScarlet-cenexin plasmid.

### siRNA transfections

h-Tert-RPE1 and h-Tert-BJ cells were transfected for 24h (*siTPX2* and *siCdk5Rap2*) 48h (*sipericentrin*) and 72h (*siKif2a*) with 20 nM siRNAs using Opti-MEM and Lipofectamine RNAiMAX (Thermofisher; 51985091 and 13778030) in MEM medium (Thermofisher; 41090) supplemented with 10% FCS and according to the manufacturer’s instructions. To deplete γ-tubulin two consecutive transfections for 72h and 48h with 40 nM siRNA were performed (Gomez-Ferreria et al., 2007). The medium was replaced 24h after transfection with DMEM supplemented with FCS 10% and 1% penicillin/streptomycin. The following sense strands of validated siRNA duplexes were used: CTRL (Qiagen, GGACCUGGAGGUCUGCUGUTT); Kif2A (Qiagen, GUUGUUUACUUUCCACGAA, Wilhelm et al., 2019); TPX2 (Qiagen, GAA UGG AAC UGG AGG GCU UTT); pericentrin (Dharmacon, UGGACGUCAUCCAAUGAGATT), Cdk5Rap2 (Dharmacon, UGGAAGAUCUCCUAACUAATT), γ-tubulin (Dharmacon, GGACAUGUUCAAGGACAACUUUGAUTT, identical sequence used in Gomez-Ferreria et al., 2007), centrobin (Dharmacon, mix of 5′-UGGAAAUGGCAGAACGAGA-3′, 5′-GCAUGAGGCUGAGCGGACA-3′, 5′-GCCCAAGAAUUGAGUCGAA-3′ and 5′-CUCCAAACCUCACGUGAUA-3′).

### Immunofluorescence

Cells were grown on glass coverslips and either fixed for 6 min at −20°C with ice-cold methanol, or with 10% formaldehyde, Pipes 20 mM pH 6.8, EGTA 10 mM, triton 0.2% for 10 min at room temperature. Cells were washed twice with phosphate-buffer saline solution (PBS) and blocked overnight at 4°C in blocking buffer (PBS, BSA 3.75%, sodium azide 0.025%). Primary antibodies were diluted in blocking buffer and incubated for 1h at room temperature. The following primary antibodies were used: cenexin (1:500, Abcam, 43840), centrin (1:2000; Merck Millipore; 04-1624; clone 20H5), centriolin (1:250, Santa Cruz biotechnology, 365521), CEP164 (1:1000; kind gift of E. Nigg, Graser et al., 2007), centrobin (1:000; Abcam; 70448),CEP192 (1:1000, Thermofisher PA5-59199), aurora T288 (1/1000, Novus biologicals, 100-2371), α-tubulin (Guerreiro and Meraldi, 2019), γ-tubulin 1/1000 (Wilhelm et al., 2019), γ-tubulin (1:1000; Sigma-Aldrich; T6557), pericentrin (1:1000, Abcam, 28144), Cdk5Rap2 (1:500, Abcam, 86340), TPX2 (1:500, Abcam, 32795), NuMa (1:1000, Abcam, 97585), Kif2A (1:500, Invitrogen, PA3 16833), katanin (1:500, Lubio Science, 14969-1-AP), TACC3 (1:500, Abcam, 134151), ch-TOG (1:200, Abcam, 236981), MCAK (1:1000, Cytoskeleton), EB1 (1:500, BD biosciences, 610535), mScarlet (1:500, Mybiosource, MBS448290). After two washes with PBS, cells were incubated with the secondary antibodies tagged with Alexa Fluor fluorophores diluted in blocking buffer (1:500) for 1h at room temperature, cells were washed twice with PBS and coverslips were mounted on slide with Vectashield medium containing DAPI (Vector laboratories). To stain cell membranes, cells were incubated (after the staining) for 15 min with Cell Mask (1:2000, Thermofisher, c10046) in the blocking buffer, rinsed and mounted with Vectashield medium containing DAPI. To stain polar chromosomes, cells were fixed in cold methanol for 6 minutes, washed and blocked in the blocking buffer. Cells expressing either GFP-centrin1 or GFP-centrin1 and mScarlet-cenexin were incubated with CENP-A antibody (1:1000, Invitrogen, MA1-20832) for 1h, washed with PBS and incubated for 1h with Alexa Fluor 488 diluted in blocking buffer (1:500). Cells were next washed twice with PBS and coverslips were mounted with Vectashield medium containing DAPI. Microscopy images were acquired using 60× and 100× (NA 1.4) oil objectives on an Olympus DeltaVision wide-field microscope (GE Healthcare) equipped with a DAPI/FITC/TRITC/Cy5 filter set (Chroma Technology Corp.) and Coolsnap HQ2 CCD camera (Roper Scientific) running Softworx (GE Healthcare). Z-stacks of 12.80-µm thickness were imaged with z-slices separated by 0.2 µm. 3D images stacks were deconvolved using Softworx (GE Healthcare) in conservative mode. Pericentrin ring images on metaphase cells were acquired using a 63x (NA 1.4) oil objective on Axio Imager LSM800 microscope (Leica) equipped with an Airyscan module. Z-slices of 0.2 µm thickness were taken to obtain a final volume containing the two centrosomes.

### Microtubule renucleation assay

h-Tert-RPE1 GFP-centrin-1 cells were treated with ice-cold medium and put on ice for 1h. Cells were either fixed 10 min with cold fixation buffer (formaldehyde 4%, Pipes 20 mM pH 6.8, EGTA 10 mM, triton 0.2%) or incubated 15s with warm DMEM medium before fixation with warm fixation buffer for 10 min. Cells were rinsed with PBS, blocked overnight at 4°C in blocking buffer (PBS, BSA 3.75%, sodium azide 0.025%), stained with anti-cenexin (1:500, Abcam 43840) and anti-α-tubulin (1:500, Guerreiro and Meraldi, 2019) primary antibodies for 1h, rinsed with PBS and incubated with secondary antibodies (Alexa Fluor 1:500) for 1h. Coverslips were mounted with Vectashield medium containing DAPI.

### Image processing and analysis

All images were processed and analysed in 3D using Imaris (Bitplane). For protein fluorescence intensity measurements at spindle poles, a sphere was drawn around the protein of interest (or based on centrin1 signal) and the mean intensity at both poles was extracted. The following sphere diameters were chosen for the quantifications (Table 1): 0.5μm for Aurora-A and Plk1, 0.65μm for CEP164, 1μm for centrobin, pericentrin, CEP192 and a 1.5μm diameter sphere was used to quantify ch-TOG, TACC3, EB1, α- and γ-tubulin fluorescence intensity at the spindle poles. The background signal was subtracted to the fluorescence intensity. The following formula was used to calculate the relative protein abundance distribution between the two poles: (pole1-pole2)/(pole1+pole2). As the localization of TPX-2, Kif2a, MCAK and katanin was not restricted to the spindle pole, the surface volume was chosen to evaluate protein concentration at both poles. For each cell, the surface volume ratio was calculated as follow: (surface1-surface2)/(surface1+surface2). To measure the half-spindle length, the distance from the pole and the center of the metaphase plate was measured for the two half-spindles (Dudka et al., 2019). For 2:2 and 1:1 cells, the spindle asymmetry index (SAI) was calculated as the ratio between the half-spindle associated to the old centrosome and the opposite half-spindle. For 1:0 cells, the half-spindle connected to the centriole-containing pole was used as reference to measure the L1 distance. For 0:0 cells, the pole and associated L1 distance were selected for the pole facing the upper side of the image. A Pearson’s correlation coefficient (or Spearman’s correlation coefficient when appropriate) was used to assess the relationship between protein distribution and associated SAI for each cell in each condition. A linear regression was performed and the value of the slope was extracted.

The relative protein distribution with respect to centrosome age was measured by taking the mean intensity within a sphere of 1 μm for pericentrin and γ-tubulin and a sphere of 2 μm for TPX2.For the daughter cell area measurements, the diameter (length+width / 2) of the two daughter cells were measured on the plane at the middle of the cell at the end of the telophase. The following formula was used to calculate the area of the two daughter cells: area = π × (diam/2)^^2^. The daughter cell area ratio was obtained by dividing the area of the daughter cell inheriting the brightest cenexin dot with the area of its sibling cell.

To evaluate the protein depletion efficiency, the sum intensity was extracted from a 1μm diameter sphere and compared to the sum intensity of the respective control condition. The relative protein distribution with respect to the centrosome was measured by taking the pole containing the brightest cenexin or centriolin dot as reference (pole1). The cortex to centrosome distances were measured on cells in metaphase and stained with the cell mask. The distances were measured on a single plane using the slice tool on Imaris. For the microtubule re-nucleation assay, the α-tubulin intensity inside a sphere of 2 μm and the mean fluorescence intensity was extracted. The pole containing the brightest cenexin was used as pole1. For the pericentrin ring images around the centrioles, a maximum intensity projection of 5-8 Z-slices was performed for each centrosome. To correlate the presence of cenexin with GFP- centrin1 brightness, the mean fluorescence intensity of both cenexin and GFP-centrin1 was extracted from a volume of 300 nm and 200x500nm respectively. All images were prepared using ImageJ/FIJI (version 2.1.0) and mounted in Inkscape (version 1.2.1).

### Statistical analysis and reproducibility

Statistical analyses were performed using GraphPad Prism9 (GraphPad). Details of the statistical tests used are indicated in the figure legends. A minimum of three independent biological replicates were performed in all experiments.

## ACKNOWLEDGEMENTS

We thank A. Khodjakov (New York State Department of Health), M.F. Tsou (National Yang-Ming University), R. Medema (Netherlands Cancer Institute) and M. Gotta (University of Geneva, Switzerland) for reagents. We thank the Flow Cytometry and the Bioimaging Facilities at the Medical Faculty of the University of Geneva for technical support, in particular N. Liaudet for his help in image data analysis. We also thank I. Gasic (University of Geneva), M. Gotta and the members of her laboratory, as well as the members of the Meraldi laboratory for helpful suggestions and critical discussions.

## Author Contributions

Conceptualization: A.T., P.M.; Formal analysis: A.T., P.M.; Investigation: A.T.; Writing – original draft: A.T.; Writing – review & editing: A.T., P.M..; Visualization: A.T.; Supervision: P.M.; Project administration: P.M.; Funding acquisition: P.M.

## SUPPLEMENTARY FIGURE LEGENDS

**Supplementary Figure 1.** (**A**) Immunofluorescence images of 2:2, 1:1, 1:0 and 0:0 RPE1 Centrin1- GFP metaphase cells stained with DAPI (blue), GFP-centrin1 (green), and Cdk5Rap2 antibody (magenta). Scale bars = 10 μm. Correlation plots between the relative Cdk5Rap2 distribution (X axis) and the SAI (Y axis) for 2:2, 1:1, 1:0 and 0:0 cells. Dots represent single cell values. The light green line indicates the slope of the linear regression. For each plot the Pearson correlation coefficient, its associated P-value, the number of cells analyzed and the slope of the linear regression are indicated. (B) Immunofluorescence images of 2:2, 1:1, 1:0 and 0:0 RPE1 GFP-centrin1 metaphase cells stained with DAPI (blue), GFP-centrin1 (green), and γ-tubulin antibody (magenta). Scale bars = 10 μm. Correlation plots between the relative γ-tubulin distribution (X axis) and the SAI (Y axis) for 2:2, 1:1, 1:0 and 0:0 cells. Dots represent single cell values. The light green line indicates the slope of the linear regression. For each plot the Pearson correlation coefficient, its associated P-value, the number of cells analyzed and the slope of the linear regression are indicated.

**Supplementary Figure 2.** (**A**) Immunofluorescence and quantification of pericentrin levels at centrosomes after *siCtrl* and *siPCNT* treatments in 1:1 RPE1 GFP-centrin1 cells. Pericentrin fluorescence intensity means of 100%, n= 72 and 3.4% ± 2.73%, n= 54 in *siCtrl* and *siPCNT* cells. (**B**) Immunofluorescence and quantification of Cdk5Rap2 levels after *siCtrl* and *siCdk5Rap2* treatments in 1:1 RPE1 GFP-centrin1 cells. Cdk5Rap2 fluorescence intensity means of 100%, n= 67 and 1.2 ± 3%, n=57 in *siCtrl* and *siCdk5Rap2* cells. (**C**) Immunofluorescence and quantification of γ-tubulin levels after *siCtrl* and *si*γ-tubulin treatment in 1:1 RPE1 GFP-centrin1 cells. γ-tubulin fluorescence intensity means of 100%, n= 54 and 8.3% ± 15.1%, n= 59 in *siCtrl* and *siγ-tubulin* cells. (**D**) Immunofluorescence and quantification of centrobin levels after *siCtrl* and *sicentrobin* treatment in 1:1 RPE1 GFP-centrin1 cells. Centrobin fluorescence intensity means of 100%, n= 23 in *siCtrl* and 5.0 ± 2.4%, n= 33 in *sicentrobin* cells.

**Supplementary Figure 3.** (**A**) Quantification of TPX2 depletion efficiency after *siCtrl* and *siTPX2* treatment in 1:1 RPE1 GFP-centrin1 cells. TPX2 fluorescence intensity means of 100%, n= 55 in *siCtrl* and 11% ± 7.9%, n= 80 in *siTPX2* cells.

**Supplementary Figure 4.** (**A**) Image sequences of parental 1:1 RPE1 GFP-centrin1 *cenexin^-/-^* cells, 1:1 RPE1 GFP-centrin1 cells expressing WT mScarlet-cenexin and 1:1 RPE1 GFP-centrin1 cells expressing S796A mScarlet-cenexin (**B**) Plots of mitotic timing (NEBD till anaphase time; NEBD = 0 min) in parental 1:1 RPE1 GFP-centrin1 *cenexin^-/-^*, 1:1 RPE1 GFP-centrin1 WT mScarlet-cenexin and 1:1 RPE1 GFP-centrin1 S796A mScarlet-cenexin cells. Median of the mitotic timing of 19.5 ± 4.9 min, n=33, 17.1 ± 2.8 min, n=76 and 20.2 ± 4.9 min respectively for the parental cell line, cells expressing mScalet-cenexin and cells expressing mScarlet-cenexin S796A. P-values = 0.30 (ns) and 0.45 (ns) for mScarlet-cenexin and mScarlet-cenexin S796A cells respectively. A Kruskal-Wallis test with Dunn’s multiple comparisons was used.

**Supplementary Figure 5.** (**A**) Immunofluorescence images of 2:2 and 1:1 RPE1 GFP-centrin1 metaphase cells stained with DAPI (blue), GFP-centrin1 (green), and cenexin antibody (yellow). Scale bars = 10 μm. Examples of 2:2 and 1:1 cells displaying the brightest centrin signal (white arrowheads) located either at the cenexin-positive centriole or at the cenexin-negative centriole. (B) Quantifications of the distribution of the brightest GFP-centrin1 signal with respect to cenexin localization. The brightest centrin signal is correlated with cenexin in only 63.6% (n= 132 cells) and 64.6% (n= 129 cells) in 2:2 and 1:1 cells respectively.

## REFERENCES

Akhmanova, A., and M.O. Steinmetz. 2019. Microtubule minus-end regulation at a glance. J Cell Sci. 132:jcs227850. doi:10.1242/jcs.227850.

Anderson, C.T., and T. Stearns. 2009. Centriole Age Underlies Asynchronous Primary Cilium Growth in Mammalian Cells. Curr Biol. 19:1498–1502. doi:10.1016/j.cub.2009.07.034.

Asteriti, I.A., F.D. Mattia, and G. Guarguaglini. 2015. Cross-Talk between AURKA and Plk1 in Mitotic Entry and Spindle Assembly. Front. Oncol. 5:283. doi:10.3389/fonc.2015.00283.

Azimzadeh, J., M.L. Wong, D.M. Downhour, A.S. Alvarado, and W.F. Marshall. 2012. Centrosome loss in the evolution of planarians. *Science (New York*, NY). 335:461–463. doi:10.1126/science.1214457.

Barr, A.R., and F. Gergely. 2007. Aurora-A: the maker and breaker of spindle poles. Journal of Cell Science. 120:2987–2996. doi:10.1242/jcs.013136.

Barr, A.R., and F. Gergely. 2008. MCAK-independent functions of ch-Tog/XMAP215 in microtubule plus-end dynamics. Molecular and cellular biology. 28:7199–7211. doi:10.1128/mcb.01040-08.

Basto, R., J. Lau, T. Vinogradova, A. Gardiol, C.G. Woods, A.L. Khodjakov, and J.W. Raff. 2006. Flies without centrioles. Cell. 125:1375–1386. doi:10.1016/j.cell.2006.05.025.

Bird, A.W., and A.A. Hyman. 2008. Building a spindle of the correct length in human cells requires the interaction between TPX2 and Aurora A. J Cell Biology. 182:289–300. doi:10.1083/jcb.200802005.

Bobinnec, Y., A.L. Khodjakov, L.M. Mir, C.L. Rieder, B. Eddé, and M. Bornens. 1998. Centriole disassembly in vivo and its effect on centrosome structure and function in vertebrate cells. J Cell Biol. 143:1575–1589.

Brunet, S., J. Dumont, K.W. Lee, K. Kinoshita, P. Hikal, O.J. Gruss, B. Maro, and M.-H. Verlhac. 2008. Meiotic Regulation of TPX2 Protein Levels Governs Cell Cycle Progression in Mouse Oocytes. PLoS ONE. 3:e3338. doi:10.1371/journal.pone.0003338.

Buffin, E., D. Emre, and R.E. Karess. 2007. Flies without a spindle checkpoint. Nature cell biology. 9:565–572. doi:10.1038/ncb1570.

Cassimeris, L., and J. Morabito. 2004. TOGp, the human homolog of XMAP215/Dis1, is required for centrosome integrity, spindle pole organization, and bipolar spindle assembly. Molecular biology of the cell. 15:1580–1590. doi:10.1091/mbc.e03-07-0544.

Chen, C., and Y.M. Yamashita. 2021. Centrosome-centric view of asymmetric stem cell division. Open Biol. 11:200314. doi:10.1098/rsob.200314.

Choi, Y.-K., P. Liu, S.K. Sze, C. Dai, and R.Z. Qi. 2010. CDK5RAP2 stimulates microtubule nucleation by the γ-tubulin ring complex. J. Cell Biol. 191:1089–1095. doi:10.1083/jcb.201007030.

Colicino, E.G., K. Stevens, E. Curtis, L. Rathbun, M. Bates, J. Manikas, J. Amack, J. Freshour, and H. Hehnly. 2019. Chromosome misalignment is associated with PLK1 activity at cenexin-positive mitotic centrosomes. Mol Biol Cell. 30:1598–1609. doi:10.1091/mbc.e18-12-0817.

Conduit, P.T., Z. Feng, J.H. Richens, J. Baumbach, A. Wainman, S.D. Bakshi, J. Dobbelaere, S. Johnson, S.M. Lea, and J.W. Raff. 2014. The Centrosome-Specific Phosphorylation of Cnn by Polo/Plk1 Drives Cnn Scaffold Assembly and Centrosome Maturation. Dev. Cell. 28:659–669. doi:10.1016/j.devcel.2014.02.013.

Conduit, P.T., A. Wainman, and J.W. Raff. 2015. Centrosome function and assembly in animal cells. Nat Rev Mol Cell Bio. 16:611–624. doi:10.1038/nrm4062.

David, A.F., P. Roudot, W.R. Legant, E. Betzig, G. Danuser, and D.W. Gerlich. 2019. Augmin accumulation on long-lived microtubules drives amplification and kinetochore-directed growth. J Cell Biol. 218:2150–2168. doi:10.1083/jcb.201805044.

Dema, A., J. van Haren, and T. Wittmann. 2022. Optogenetic EB1 inactivation shortens metaphase spindles by disrupting cortical force-producing interactions with astral microtubules. Curr. Biol. 32:1197–1205.e4. doi:10.1016/j.cub.2022.01.017.

Dictenberg, J.B., W. Zimmerman, C.A. Sparks, A. Young, C. Vidair, Y. Zheng, W. Carrington, F.S. Fay, and S.J. Doxsey. 1998. Pericentrin and γ-Tubulin Form a Protein Complex and Are Organized into a Novel Lattice at the Centrosome. J. Cell Biol. 141:163–174. doi:10.1083/jcb.141.1.163.

Domnitz, S.B., M. Wagenbach, J. Decarreau, and L. Wordeman. 2012. MCAK activity at microtubule tips regulates spindle microtubule length to promote robust kinetochore attachment. J. Cell Biol. 197:231–237. doi:10.1083/jcb.201108147.

Dudka, D., C. Castrogiovanni, N. Liaudet, H. Vassal, and P. Meraldi. 2019. Spindle-Length-Dependent HURP Localization Allows Centrosomes to Control Kinetochore-Fiber Plus-End Dynamics. Curr Biol. 29:3563–3578.e6. doi:10.1016/j.cub.2019.08.061.

Dudka, D., and P. Meraldi. 2017. Asymmetric Cell Division in Development, Differentiation and Cancer. Results Problems Cell Differ. 61:301–321. doi:10.1007/978-3-319-53150-2_14.

Dumont, S., and T.J. Mitchison. 2009. Force and length in the mitotic spindle. Current biology : CB. 19:R749–61. doi:10.1016/j.cub.2009.07.028.

Fırat-Karalar, E.N., and T. Stearns. 2014. The centriole duplication cycle. Philos. Trans. R. Soc. B: Biol. Sci. 369:20130460. doi:10.1098/rstb.2013.0460.

Fong, K.-W., Y.-K. Choi, J.B. Rattner, and R.Z. Qi. 2008. CDK5RAP2 Is a Pericentriolar Protein That Functions in Centrosomal Attachment of the γ-Tubulin Ring Complex. Mol Biol Cell. 19:115–125. doi:10.1091/mbc.e07-04-0371.

Fu, J., M. Bian, G. Xin, Z. Deng, J. Luo, X. Guo, H. Chen, Y. Wang, Q. Jiang, and C. Zhang. 2015. TPX2 phosphorylation maintains metaphase spindle length by regulating microtubule flux. J Cell Biol. 210:373–383. doi:10.1083/jcb.201412109.

Fuse, N., K. Hisata, A.L. Katzen, and F. Matsuzaki. 2003. Heterotrimeric G Proteins Regulate Daughter Cell Size Asymmetry in Drosophila Neuroblast Divisions. Curr Biol. 13:947–954. doi:10.1016/s0960-9822(03)00334-8.

Gaetz, J., and T.M. Kapoor. 2004. Dynein/dynactin regulate metaphase spindle length by targeting depolymerizing activities to spindle poles. J Cell Biol. 166:465–471. doi:10.1083/jcb.200404015.

Ganem, N.J., and D.A. Compton. 2004. The KinI kinesin Kif2a is required for bipolar spindle assembly through a functional relationship with MCAK. J Cell Biol. 166:473–478. doi:10.1083/jcb.200404012.

Gasic, I., P. Nerurkar, and P. Meraldi. 2015. Centrosome age regulates kinetochore microtubule stability and biases chromosome mis-segregation. eLife. 4:e07909. doi:10.7554/elife.07909.

Gergely, F., V.M. Draviam, and J.W. Raff. 2003. The ch-TOG/XMAP215 protein is essential for spindle pole organization in human somatic cells. Genes Dev. 17:336–341. doi:10.1101/gad.245603.

Goshima, G., M. Mayer, N. Zhang, N. Stuurman, and R.D. Vale. 2008. Augmin: a protein complex required for centrosome-independent microtubule generation within the spindle. J Cell Biol. 181:421–429. doi:10.1083/jcb.200711053.

Goshima, G., and J.M. Scholey. 2010. Control of mitotic spindle length. Annual Review of Cell and Developmental Biology. 26:21–57. doi:10.1146/annurev-cellbio-100109-104006.

Goshima, G., R. Wollman, N. Stuurman, J.M. Scholey, and R.D. Vale. 2005. Length control of the metaphase spindle. Current biology : CB. 15:1979–1988. doi:10.1016/j.cub.2005.09.054.

Greenan, G., C.P. Brangwynne, S. Jaensch, J. Gharakhani, F. Jülicher, and A.A. Hyman. 2010. Centrosome size sets mitotic spindle length in Caenorhabditis elegans embryos. Current biology : CB. 20:353–358. doi:10.1016/j.cub.2009.12.050.

Gruss, O.J., M. Wittmann, H. Yokoyama, R. Pepperkok, T. Kufer, H. Silljé, E. Karsenti, I.W. Mattaj, and I. Vernos. 2002. Chromosome-induced microtubule assembly mediated by TPX2 is required for spindle formation in HeLa cells. Nat. Cell Biol. 4:871–879. doi:10.1038/ncb870.

Guerreiro, A., F.D. Sousa, N. Liaudet, D. Ivanova, A. Eskat, and P. Meraldi. 2021. WDR62 localizes katanin at spindle poles to ensure synchronous chromosome segregation. J Cell Biol. 220:e202007171. doi:10.1083/jcb.202007171.

Hayward, D., J. Metz, C. Pellacani, and J.G. Wakefield. 2014. Synergy between multiple microtubule-generating pathways confers robustness to centrosome-driven mitotic spindle formation. Developmental cell. 28:81–93. doi:10.1016/j.devcel.2013.12.001.

Heald, R., R. Tournebize, T. Blank, R. Sandaltzopoulos, P. Becker, A.A. Hyman, and E. Karsenti. 1996. Self-organization of microtubules into bipolar spindles around artificial chromosomes in Xenopus egg extracts. Nature. 382:420–425. doi:10.1038/382420a0.

Helmke, K.J., and R. Heald. 2014. TPX2 levels modulate meiotic spindle size and architecture in Xenopus egg extracts. J. cell Biol. 206:385–93. doi:10.1083/jcb.201401014.

Huang, J., Z. Liang, C. Guan, S. Hua, and K. Jiang. 2021. WDR62 regulates spindle dynamics as an adaptor protein between TPX2/Aurora A and katanin. J Cell Biology. 220:e202007167. doi:10.1083/jcb.202007167.

Ishikawa, H., A. Kubo, S. Tsukita, and S. Tsukita. 2005. Odf2-deficient mother centrioles lack distal/subdistal appendages and the ability to generate primary cilia. Nature cell biology. 7:517–524. doi:10.1038/ncb1251.

Jang, C.-Y., J.A. Coppinger, A. Seki, J.R. Yates, and G. Fang. 2009. Plk1 and Aurora A regulate the depolymerase activity and the cellular localization of Kif2a. Journal of Cell Science. 122:1334– 1341. doi:10.1242/jcs.044321.

Januschke, J., J. Reina, S. Llamazares, T. Bertran, F. Rossi, J. Roig, and C. Gonzalez. 2013. Centrobin controls mother-daughter centriole asymmetry in Drosophila neuroblasts. Nature cell biology. 15:241–248. doi:10.1038/ncb2671.

Jeffery, J.M., A.J. Urquhart, V.N. Subramaniam, R.G. Parton, and K.K. Khanna. 2010. Centrobin regulates the assembly of functional mitotic spindles. Oncogene. 29:2649–2658. doi:10.1038/onc.2010.37.

Joukov, V., J.C. Walter, and A. De Nicolo. 2014. The Cep192-Organized Aurora A-Plk1 Cascade Is Essential for Centrosome Cycle and Bipolar Spindle Assembly. Mol Cell. 55:578–591. doi:10.1016/j.molcel.2014.06.016.

Keller, L.C., K.A. Wemmer, and W.F. Marshall. 2010. Influence of centriole number on mitotic spindle length and symmetry. *Cytoskeleton (Hoboken*, N.J.). 67:504–518. doi:10.1002/cm.20462.

Khodjakov, A.L., R.W. Cole, B.R. Oakley, and C.L. Rieder. 2000. Centrosome-independent mitotic spindle formation in vertebrates. Current biology : CB. 10:59–67.

Khodjakov, A.L., and C.L. Rieder. 2001. Centrosomes enhance the fidelity of cytokinesis in vertebrates and are required for cell cycle progression. J Cell Biol. 153:237–242.

Kiyomitsu, T., and I.M. Cheeseman. 2013. Cortical Dynein and asymmetric membrane elongation coordinately position the spindle in anaphase. Cell. 154:391–402. doi:10.1016/j.cell.2013.06.010.

Kong, D., V. Farmer, A. Shukla, J. James, R. Gruskin, S. Kiriyama, and J. Loncarek. 2014. Centriole maturation requires regulated Plk1 activity during two consecutive cell cycles. J. Cell Biol. 206:855–865. doi:10.1083/jcb.201407087.

Kong, D., N. Sahabandu, C. Sullenberger, A. Vásquez-Limeta, D. Luvsanjav, K. Lukasik, and J. Loncarek. 2020. Prolonged mitosis results in structurally aberrant and over-elongated centrioles. J. Cell Biol. 219:e201910019. doi:10.1083/jcb.201910019.

Kufer, T.A., H.H.W. Sillje, R. Körner, O.J. Gruss, P. Meraldi, and E.A. Nigg. 2002. Human TPX2 is required for targeting Aurora-A kinase to the spindle. J Cell Biol. 158:617–623. doi:10.1083/jcb.200204155.

Lacroix, B., and J. Dumont. 2022. Spatial and Temporal Scaling of Microtubules and Mitotic Spindles. Cells. 11:248. doi:10.3390/cells11020248.

Lane, H.A., and E.A. Nigg. 1996. Antibody microinjection reveals an essential role for human polo-like kinase 1 (Plk1) in the functional maturation of mitotic centrosomes. J. cell Biol. 135:1701– 1713. doi:10.1083/jcb.135.6.1701.

Lawo, S., M. Hasegan, G.D. Gupta, and L. Pelletier. 2012. Subdiffraction imaging of centrosomes reveals higher-order organizational features of pericentriolar material. Nat Cell Biol. 14:1148– 1158. doi:10.1038/ncb2591.

Lee, K., and K. Rhee. 2011. PLK1 phosphorylation of pericentrin initiates centrosome maturation at the onset of mitosis. J. Cell Biol. 195:1093–1101. doi:10.1083/jcb.201106093.

Logarinho, E., S. Maffini, M. Barisic, A. Marques, A. Toso, P. Meraldi, and H. Maiato. 2012. CLASPs prevent irreversible multipolarity by ensuring spindle-pole resistance to traction forces during chromosome alignment. Nature cell biology. 14:295–303. doi:10.1038/ncb2423.

Loughlin, R., J.D. Wilbur, F.J. McNally, F.J. Nédélec, and R. Heald. 2011. Katanin contributes to interspecies spindle length scaling in Xenopus. Cell. 147:1397–1407. doi:10.1016/j.cell.2011.11.014.

Maiato, H., A.L. Khodjakov, and C.L. Rieder. 2005. Drosophila CLASP is required for the incorporation of microtubule subunits into fluxing kinetochore fibres. Nature cell biology. 7:42–47. doi:10.1038/ncb1207.

Mayr, M.I., S. Hümmer, J. Bormann, T. Grüner, S. Adio, G. Woehlke, and T.U. Mayer. 2007. The human kinesin Kif18A is a motile microtubule depolymerase essential for chromosome congression. Current biology : CB. 17:488–498. doi:10.1016/j.cub.2007.02.036.

Mazo, G., N. Soplop, W.-J. Wang, K. Uryu, and M.-F.B. Tsou. 2016. Spatial Control of Primary Ciliogenesis by Subdistal Appendages Alters Sensation-Associated Properties of Cilia. Dev Cell. 39:424–437. doi:10.1016/j.devcel.2016.10.006.

McNally, K., A. Audhya, K. Oegema, and F.J. McNally. 2006. Katanin controls mitotic and meiotic spindle length. J. Cell Biol. 175:881–891. doi:10.1083/jcb.200608117.

Mennella, V., B. Keszthelyi, K.L. McDonald, B. Chhun, F. Kan, G.C. Rogers, B. Huang, and D.A. Agard. 2012. Subdiffraction-resolution fluorescence microscopy reveals a domain of the centrosome critical for pericentriolar material organization. Nat. Cell Biol. 14:1159–1168. doi:10.1038/ncb2597.

Meraldi, P. 2016. Centrosomes in spindle organization and chromosome segregation: a mechanistic view. Chromosome Res. 24:19–34. doi:10.1007/s10577-015-9508-2.

Miller, K.E., A.M. Session, and R. Heald. 2019. Kif2a Scales Meiotic Spindle Size in Hymenochirus boettgeri. Curr. Biol. 29:3720–3727.e5. doi:10.1016/j.cub.2019.08.073.

Mogessie, B., K. Scheffler, and M. Schuh. 2018. Assembly and Positioning of the Oocyte Meiotic Spindle. Annu. Rev. Cell Dev. Biol. 34:381–403. doi:10.1146/annurev-cellbio-100616-060553.

Moritz, M., M.B. Braunfeld, J.W. Sedat, B. Alberts, and D.A. Agard. 1995. Microtubule nucleation by γ-tubulin-containing rings in the centrosome. Nature. 378:638–640. doi:10.1038/378638a0.

Nigg, E.A., and A.J. Holland. 2018. Once and only once: mechanisms of centriole duplication and their deregulation in disease. Nat. Rev. Mol. Cell Biol. 19:297–312. doi:10.1038/nrm.2017.127.

Nigg, E.A., and T. Stearns. 2011. The centrosome cycle: Centriole biogenesis, duplication and inherent asymmetries. Nature cell biology. 13:1154–1160. doi:10.1038/ncb2345.

Ohta, M., Z. Zhao, D. Wu, S. Wang, J.L. Harrison, J.S. Gómez-Cavazos, A. Desai, and K.F. Oegema. 2021. Polo-like kinase 1 independently controls microtubule-nucleating capacity and size of the centrosome. J Cell Biology. 220:e202009083. doi:10.1083/jcb.202009083.

Palazzo, R.E., J.M. Vogel, B.J. Schnackenberg, D.R. Hull, and X. Wu. 2000. Centrosome maturation. Current topics in developmental biology. 49:449–470.

Pereira, G., T.U. Tanaka, K. Nasmyth, and E. Schiebel. 2001. Modes of spindle pole body inheritance and segregation of the Bfa1p–Bub2p checkpoint protein complex. EMBO J. 20:6359–6370. doi:10.1093/emboj/20.22.6359.

Petry, S. 2016. Mechanisms of Mitotic Spindle Assembly. Annu Rev Biochem. 85:659–83. doi:10.1146/annurev-biochem-060815-014528.

Petry, S., A.C. Groen, K. Ishihara, T.J. Mitchison, and R.D. Vale. 2013. Branching Microtubule Nucleation in Xenopus Egg Extracts Mediated by Augmin and TPX2. Cell. 152:768–777. doi:10.1016/j.cell.2012.12.044.

Polverino, F., F.D. Naso, I.A. Asteriti, V. Palmerini, D. Singh, D. Valente, A.W. Bird, A. Rosa, M. Mapelli, and G. Guarguaglini. 2021. The Aurora-A/TPX2 Axis Directs Spindle Orientation in Adherent Human Cells by Regulating NuMA and Microtubule Stability. Curr. Biol. 31:658–667.e5. doi:10.1016/j.cub.2020.10.096.

Rathbun, L.I., A.A. Aljiboury, X. Bai, N.A. Hall, J. Manikas, J.D. Amack, J.N. Bembenek, and H. Hehnly. 2020. PLK1-and PLK4-Mediated Asymmetric Mitotic Centrosome Size and Positioning in the Early Zebrafish Embryo. Curr. Biol. 30:4519–4527.e3. doi:10.1016/j.cub.2020.08.074.

Reber, S.B., J. Baumgart, P.O. Widlund, A. Pozniakovsky, J. Howard, A.A. Hyman, and F. Jülicher. 2013. XMAP215 activity sets spindle length by controlling the total mass of spindle microtubules. Nature cell biology. 15:1116–1122. doi:10.1038/ncb2834.

Ren, X., and D.A. Weisblat. 2006. Asymmetrization of first cleavage by transient disassembly of one spindle pole aster in the leech Helobdella robusta. Dev. Biol. 292:103–115. doi:10.1016/j.ydbio.2005.12.049.

Roubinet, C., A. Tsankova, T.T. Pham, A. Monnard, E. Caussinus, M. Affolter, and C. Cabernard. 2017. Spatio-temporally separated cortical flows and spindle geometry establish physical asymmetry in fly neural stem cells. Nat Commun. 8:1383. doi:10.1038/s41467-017-01391-w.

Roux-Bourdieu, M.L., D. Dwivedi, D. Harry, and P. Meraldi. 2022. PLK1 controls centriole distal appendage formation and centrobin removal via independent pathways. J Cell Sci. 135. doi:10.1242/jcs.259120.

Sanchez, A.D., and J.L. Feldman. 2017. Microtubule-organizing centers: from the centrosome to non-centrosomal sites. Curr. Opin. Cell Biol. 44:93–101. doi:10.1016/j.ceb.2016.09.003.

Shcheprova, Z., S. Baldi, S.B. Frei, G. Gonnet, and Y. Barral. 2008. A mechanism for asymmetric segregation of age during yeast budding. Nature. 454:728–734. doi:10.1038/nature07212.

Sir, J.-H., M. Pütz, O. Daly, C.G. Morrison, M. Dunning, J.V. Kilmartin, and F. Gergely. 2013. Loss of centrioles causes chromosomal instability in vertebrate somatic cells. J Cell Biol. 203:747–756. doi:10.1083/jcb.201309038.

Sobajima, T., K.M. Kowalczyk, S. Skylakakis, D. Hayward, L.J. Fulcher, C. Neary, C. Batley, S. Kurlekar, E. Roberts, U. Gruneberg, and F.A. Barr. 2023. PP6 regulation of Aurora A–TPX2 limits NDC80 phosphorylation and mitotic spindle size. J Cell Biology. 222:e202205117. doi:10.1083/jcb.202205117.

Soung, N.-K., J.-E. Park, L.-R. Yu, K.H. Lee, J.-M. Lee, J.K. Bang, T.D. Veenstra, K. Rhee, and K.S. Lee. 2009. Plk1-dependent and -independent roles of an ODF2 splice variant, hCenexin1, at the centrosome of somatic cells. Developmental cell. 16:539–550. doi:10.1016/j.devcel.2009.02.004.

Sullenberger, C., A. Vasquez-Limeta, D. Kong, and J. Loncarek. 2020. With Age Comes Maturity: Biochemical and Structural Transformation of a Human Centriole in the Making. Cells. 9:1429. doi:10.3390/cells9061429.

Tan, C.H., J. Pines, I. Gasic, S.P. Huber-Reggi, D. Dudka, M. Barisic, H. Maiato, and P. Meraldi. 2015. The equatorial position of the metaphase plate ensures symmetric cell divisions. eLife. 4:e05124. doi:10.7554/elife.05124.

Uehara, R., R. Nozawa, A. Tomioka, S. Petry, R.D. Vale, C. Obuse, and G. Goshima. 2009. The augmin complex plays a critical role in spindle microtubule generation for mitotic progression and cytokinesis in human cells. Proc. Natl. Acad. Sci. 106:6998–7003. doi:10.1073/pnas.0901587106.

Vasquez-Limeta, A., and J. Loncarek. 2021. Human centrosome organization and function in interphase and mitosis. Semin. cell Dev. Biol. 117:30–41. doi:10.1016/j.semcdb.2021.03.020.

Wang, X., J.-W. Tsai, J.H. Imai, W.-N. Lian, R.B. Vallee, and S.-H. Shi. 2009. Asymmetric centrosome inheritance maintains neural progenitors in the neocortex. Nature. 461:947–955. doi:10.1038/nature08435.

Watanabe, S., F. Meitinger, A.K. Shiau, K. Oegema, and A. Desai. 2020. Centriole-independent mitotic spindle assembly relies on the PCNT–CDK5RAP2 pericentriolar matrix. J Cell Biology. 219:e202006010. doi:10.1083/jcb.202006010.

Wilhelm, T., A.-M. Olziersky, D. Harry, F.D. Sousa, H. Vassal, A. Eskat, and P. Meraldi. 2019. Mild replication stress causes chromosome mis-segregation via premature centriole disengagement. Nat Commun. 10:3585. doi:10.1038/s41467-019-11584-0.

Wong, Y.L., J.V. Anzola, R.L. Davis, M. Yoon, A. Motamedi, A. Kroll, C.P. Seo, J.E. Hsia, S.K. Kim, J.W. Mitchell, B.J. Mitchell, A.B. Desai, T.C. Gahman, A.K. Shiau, and K. Oegema. 2015. Cell biology. Reversible centriole depletion with an inhibitor of Polo-like kinase 4. Science (New York, NY). 348:1155–1160. doi:10.1126/science.aaa5111.

Wu, J., A. Larreategui-Aparicio, M.L.A. Lambers, D.L. Bodor, S.J. Klaasen, E. Tollenaar, M. de Ruijter-Villani, and G.J.P.L. Kops. 2023. Microtubule nucleation from the fibrous corona by LIC1-pericentrin promotes chromosome congression. Curr Biol. 33:912–925.e6. doi:10.1016/j.cub.2023.01.010.

Wu, T., J. Dong, J. Fu, Y. Kuang, B. Chen, H. Gu, Y. Luo, R. Gu, M. Zhang, W. Li, X. Dong, X. Sun, Q. Sang, and L. Wang. 2022. The mechanism of acentrosomal spindle assembly in human oocytes. Science. 378:eabq7361. doi:10.1126/science.abq7361.

Yamashita, Y.M., A.P. Mahowald, J.R. Perlin, and M.T. Fuller. 2007. Asymmetric inheritance of mother versus daughter centrosome in stem cell division. *Science (New York*, NY). 315:518–521. doi:10.1126/science.1134910.

Yi, P., and G. Goshima. 2018. Microtubule nucleation and organization without centrosomes. Curr. Opin. plant Biol. 46:1–7. doi:10.1016/j.pbi.2018.06.004.

Zou, C., J. Li, Y. Bai, W.T. Gunning, D.E. Wazer, V. Band, and Q. Gao. 2005. Centrobin: a novel daughter centriole-associated protein that is required for centriole duplication. J Cell Biol. 171:437–445. doi:10.1083/jcb.200506185.

