## Supplemental Figures for "Centrosome age breaks spindle size symmetry even in “symmetrically” dividing cells"

### Thomas and Meraldi - Supplementary figure 1

A

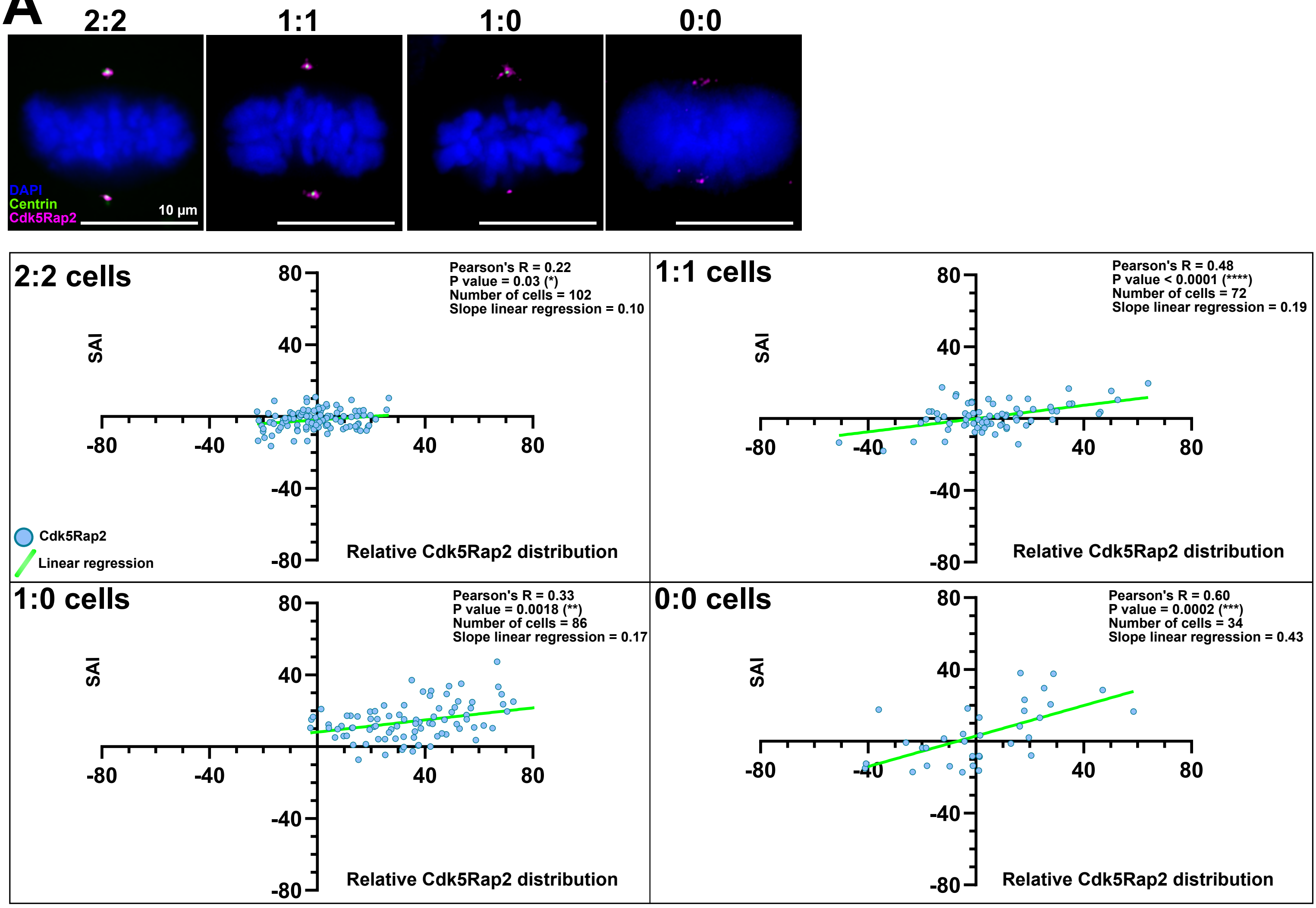


B

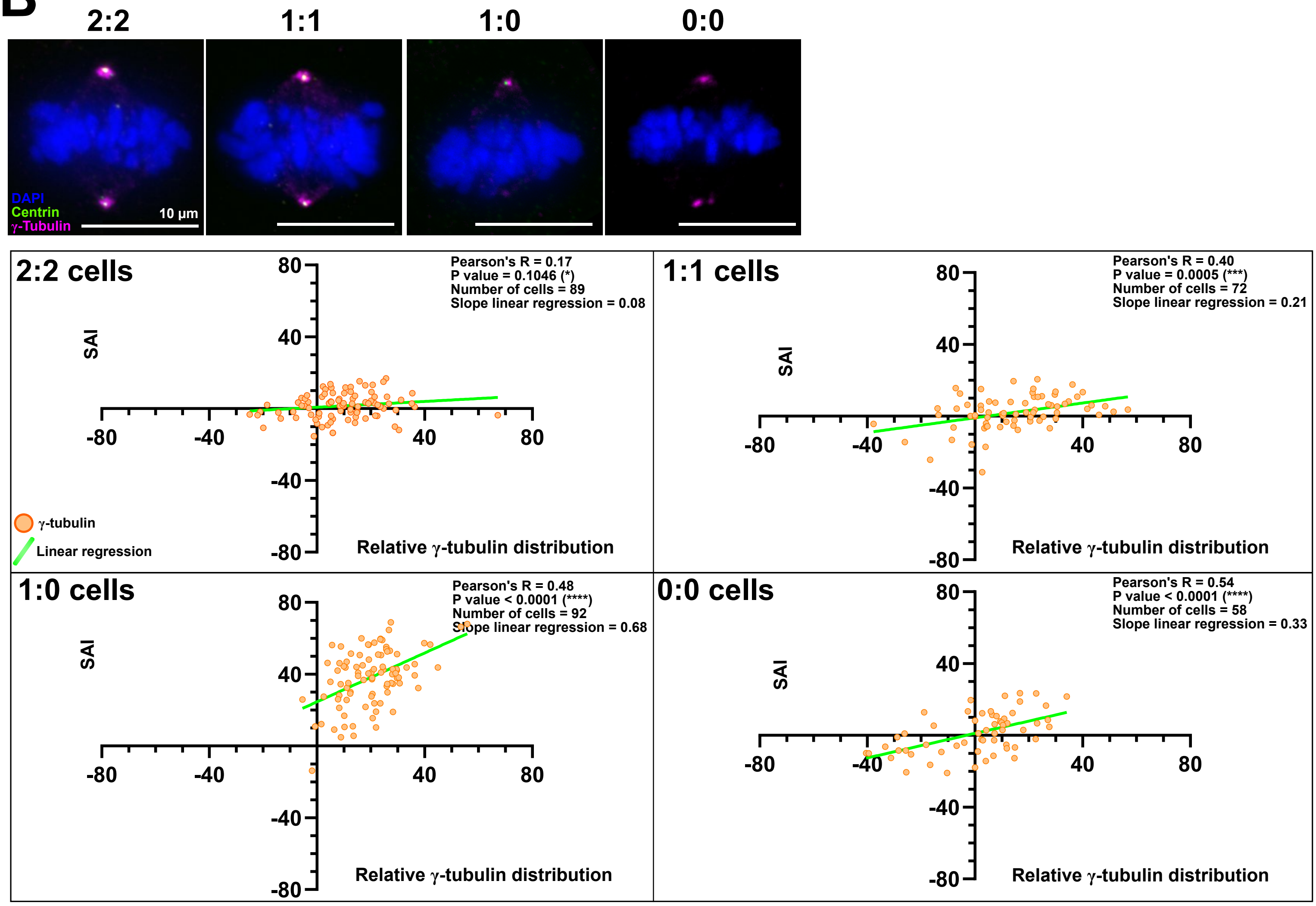

### Thomas and Meraldi - Supplementary figure 2

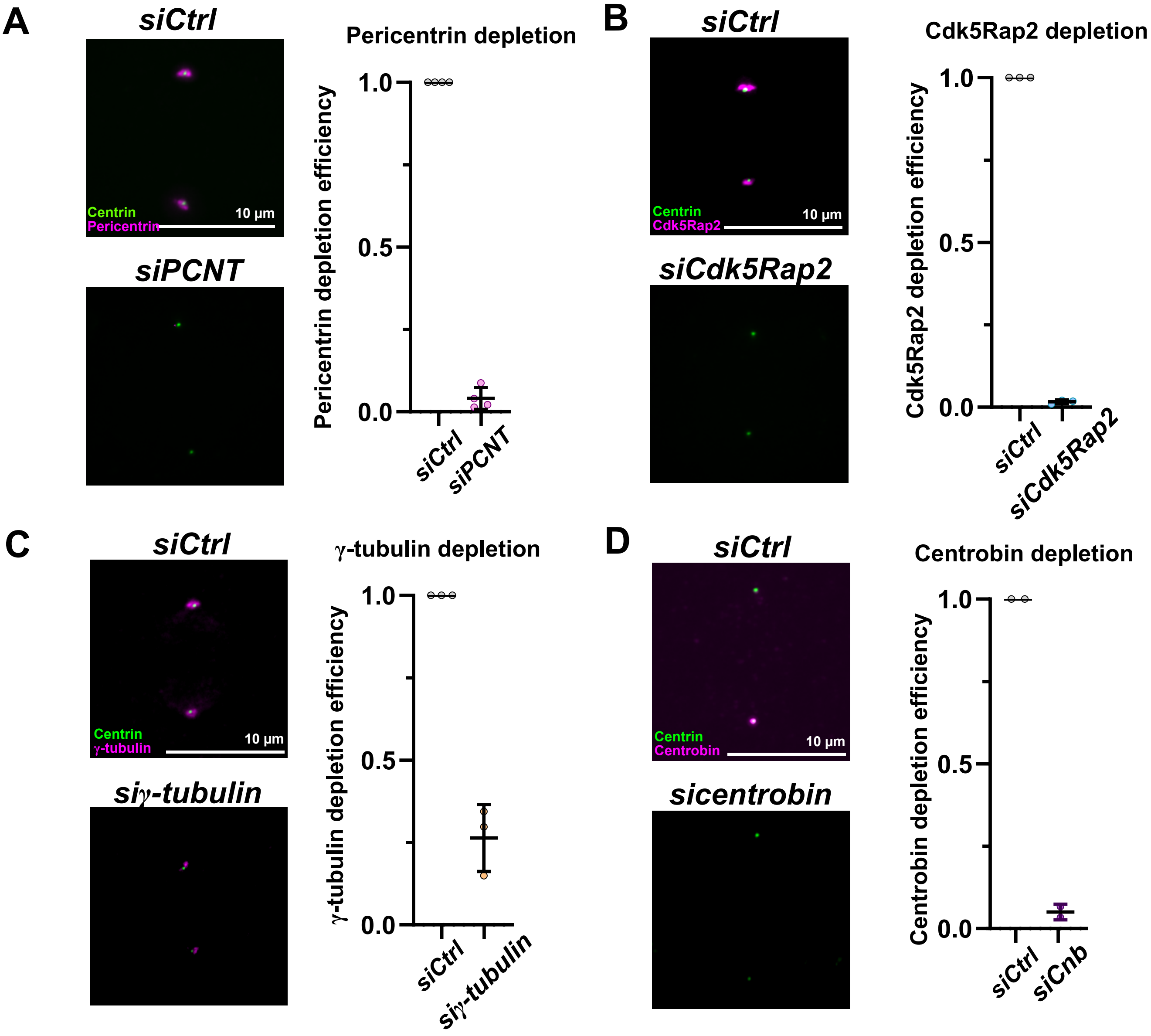

### Thomas and Meraldi - Supplementary figure 3

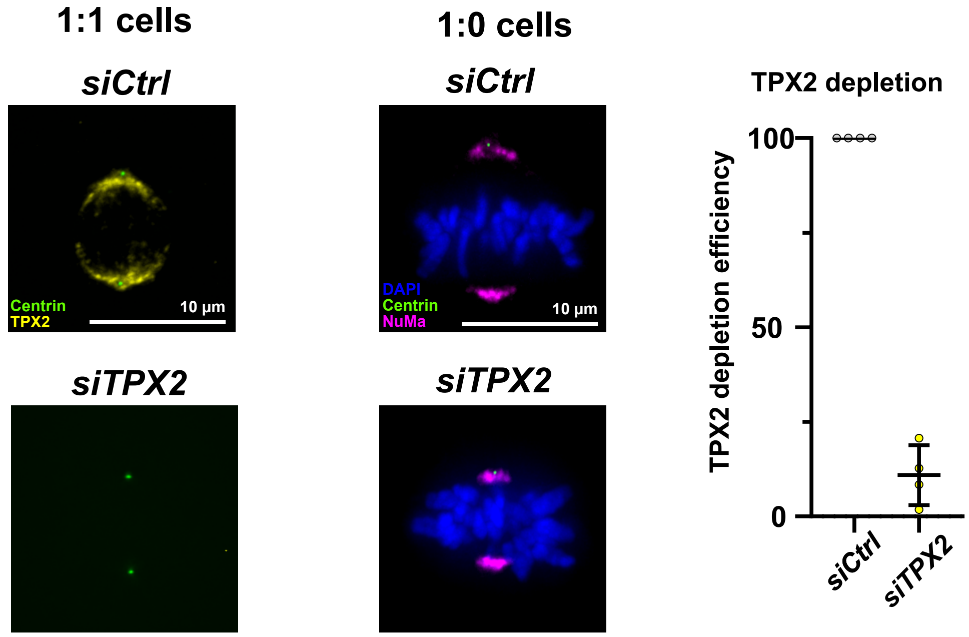

### Thomas and Meraldi - Supplementary figure 4

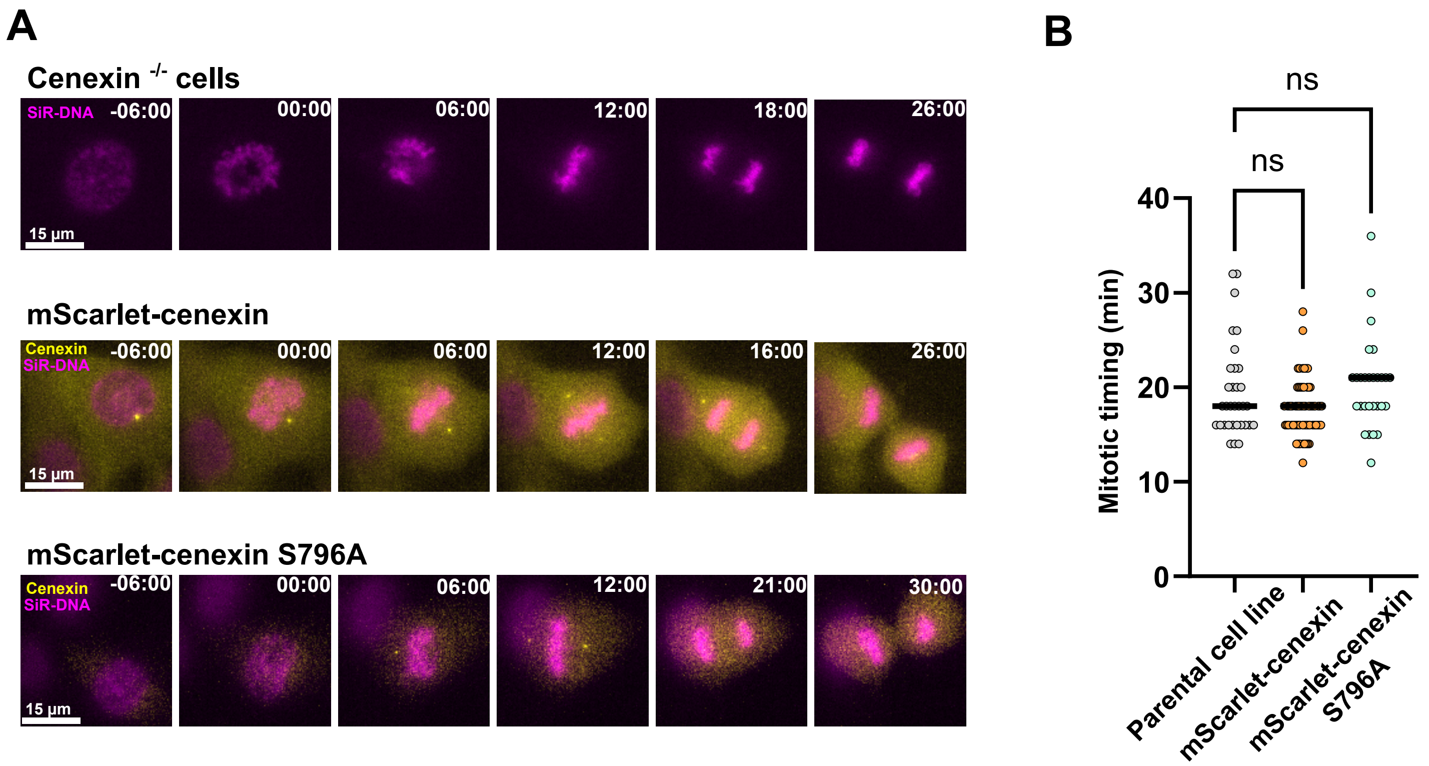

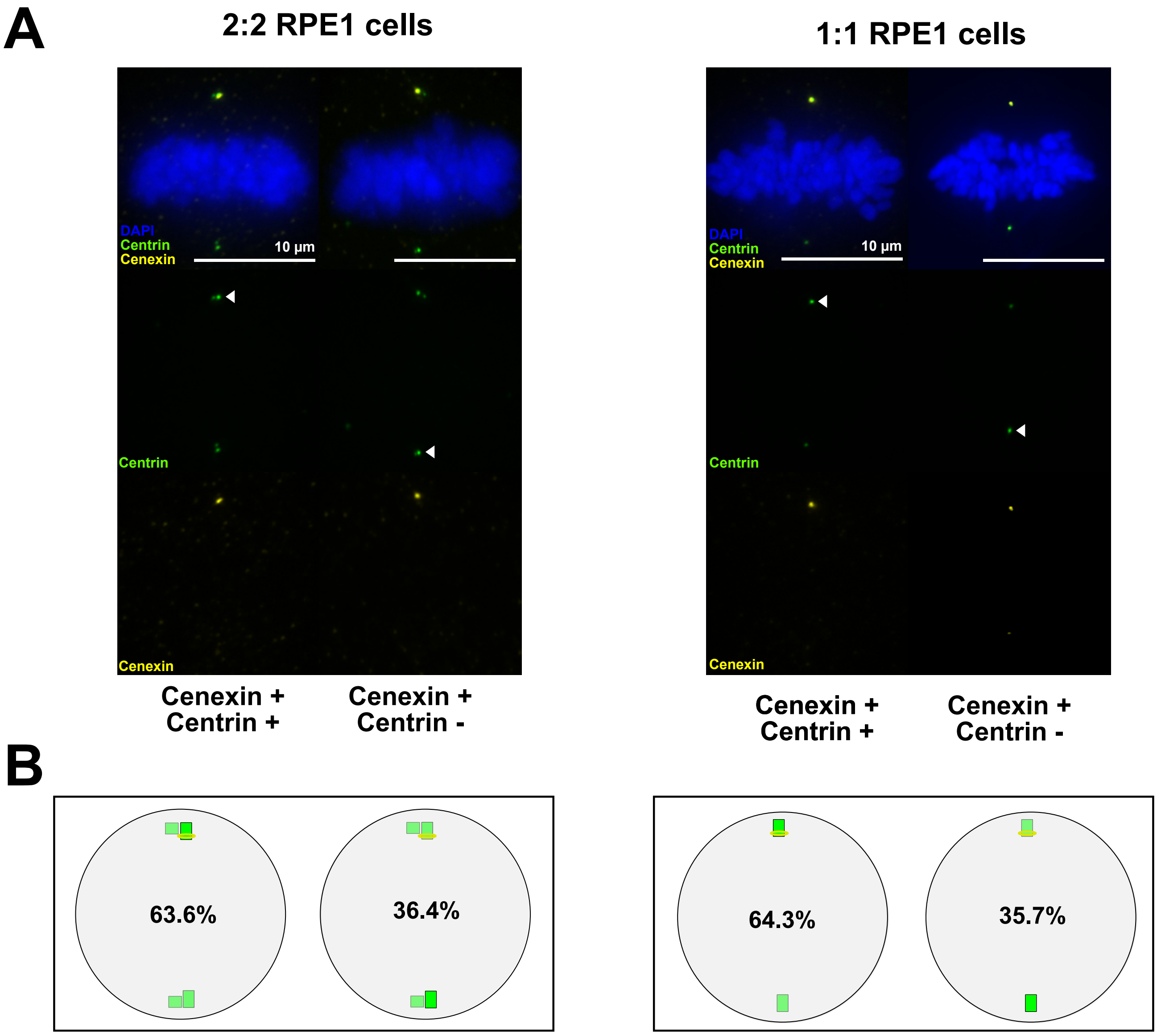
A
2:2 RPE1 cells
1:1 RPE1 cells
DAPI
Centrin
Cenexin
10 µm
Centrin
Cenexin
DAPI
Centrin
Cenexin
10 µm
Centrin
Cenexin
Cenexin +
Centrin +
Cenexin +
Centrin -
Cenexin +
Centrin +
Cenexin +
Centrin -
B
63.6%
36.4%
64.3%
35.7%
